# Prefrontal responses during proactive and reactive inhibition are differentially impacted by stress in anorexia and bulimia nervosa

**DOI:** 10.1101/2020.02.27.968719

**Authors:** Margaret L. Westwater, Flavia Mancini, Adam X. Gorka, Jane Shapleske, Jaco Serfontein, Christian Grillon, Monique Ernst, Hisham Ziauddeen, Paul C. Fletcher

**Affiliations:** Department of Psychiatry, University of Cambridge, Herchel Smith Building, Addenbrooke’s Hospital, Cambridge CB2 0SZ, UK; National Institute of Mental Health, National Institutes of Health, Bethesda, MD, USA; Computational and Biological Learning Laboratory, Department of Engineering, University of Cambridge, Cambridge CB2 1PZ; Adult Eating Disorders Service, Cambridgeshire and Peterborough NHS Foundation Trust, Cambridge, CB21 5EF; Wellcome Trust MRC Institute of Metabolic Science, University of Cambridge, Cambridge Biomedical Campus, Cambridge CB2 0QQ, UK

**Keywords:** inhibitory control, stress, eating disorders, binge-eating, fMRI, vmPFC

## Abstract

**Background:** Binge-eating is a distressing, transdiagnostic eating disorder symptom associated with impulsivity, particularly in negative mood states. Neuroimaging studies of bulimia nervosa (BN) report reduced activity in fronto-striatal regions implicated in self-regulatory control. However, it remains unknown if negative affective states, including stress, impair self-regulation, and, if so, whether such self-regulatory deficits generalize to binge-eating in underweight individuals (i.e., the bingeing/purging subtype of anorexia nervosa; AN-BP).

**Methods:** We determined the effect of acute stress on inhibitory control in 85 women (33 BN, 22 AN-BP, 30 matched controls). Participants underwent repeated functional MRI scanning, during performance of the stop-signal anticipation task, a validated measure of proactive (i.e., anticipation of stopping) and reactive (outright stopping) inhibition. Neural and behavioral responses to induced, psychological stress and a control task were evaluated on two separate days.

**Results:** Women with BN had reduced proactive inhibition while prefrontal responses were increased in both AN-BP and BN. Reactive inhibition was neurally and behaviorally intact in both diagnostic groups. Both AN-BP and BN groups showed distinct, stress-induced changes in prefrontal activity during both proactive and reactive inhibition. However, task performance was not significantly affected by stress.

**Conclusions:** These findings offer novel evidence of reduced proactive inhibition in BN, yet inhibitory control deficits did not generalize to AN-BP. While both groups showed altered neural responses during inhibition following stress, neither group demonstrated stress-induced performance deficits. As such, our findings counsel against a simplistic, stress-induced failure of regulation as a holistic explanation for binge-eating in these conditions.

## 1. Introduction

Anorexia Nervosa (AN) and Bulimia Nervosa (BN) are complex psychiatric conditions marked by abnormal eating behavior and distorted thoughts surrounding food, eating and body perception. They share cardinal symptoms, including recurrent binge-eating and compensatory behaviors (e.g., vomiting, laxative use), which occur in both BN and the binge-eating and purging subtype of AN (AN-BP; 1). Binge-eating engenders substantial distress, and it is associated with significant impairment and comorbidity (2, 3). Although this syndrome has been related to aberrant reward and self-regulatory processing (4–6), its pathophysiological correlates remain poorly characterized, particularly in the severely understudied condition, AN-BP. As binge-eating has been predominantly studied in BN and binge-eating disorder, it remains unclear if models of binge-eating based on these conditions generalize to AN-BP, where individuals endure significant weight loss.

An influential model of binge-eating posits that it emerges in response to negative affective states, which reduce an individual’s capacity for self-control, thereby leading to loss-of-control eating (7). While elevated trait impulsivity in BN (8) and, to a lesser extent AN-BP (9), lends some support to this model, experimental studies of self-regulation are more equivocal due to inconsistencies across neural and behavioral findings. For example, functional magnetic resonance imaging (fMRI) studies of adolescent (10) and adult (11, 12) BN report reduced fronto-striatal activity during conflict and action inhibition trials on Simon Spatial and Go/NoGo tasks, respectively, yet behavioral impairments were only observed on the Simon Spatial task in adult BN. Similarly, adolescents with mixed binge-purge pathology have shown increased bilateral hypothalamic, precentral gyrus and right dorsolateral prefrontal cortex (dlPFC), anterior cingulate cortex (ACC) and middle temporal gyrus activity relative to controls (13) during intact Go/NoGo performance. Altered brain activity in the absence of behavioral impairment could indicate either inefficient or compensatory neural responses to preserve task performance. Interestingly, despite nonsignificant differences in stop-signal performance at baseline, augmented medial prefrontal and ACC activity on failed stop-signal trials predicted the onset of eating disorder (ED) behaviors at two-year, longitudinal follow-up (14).

Inconsistencies across levels of analysis and cognitive tasks could partly reflect heterogeneity within the theoretical construct of ‘self-control.’ Behavioral (15–17) and neurobiological data support dissociable forms of impulsivity, including ‘temporal impulsivity’, relating to the delayed receipt of reward, and ‘response inhibition’, or ‘inhibitory control’ (IC), which is the capacity to slow or stop a response tendency (18). Moreover, theoretical frameworks suggest that IC is modulated by both proactive and reactive processes (19, 20). ‘Proactive inhibition’ describes a goal-directed process, elicited by predictive cues, which is used to restrain actions in preparation for stopping. In contrast, ‘reactive inhibition’ is a stimulus-driven process, where a salient signal triggers action cancellation. These inhibitory modes have both shared and unique neural correlates (21). Bilateral frontoparietal and basal ganglia regions form a broad IC network that subserves both processes, but bilateral superior parietal and right-dominant, frontal, temporal and parietal regions have been uniquely related to proactive and reactive inhibition, respectively (22, 23). As such, the neural and behavioral distinctions between proactive and reactive inhibition should be considered when attempting to identify the specific self-regulatory impairment(s) associated with binge-eating disorders.

In addition to examining specific self-regulatory deficits in AN-BP and BN, efforts to validate the model must also consider the impact of negative mood states on this process. While experience sampling has shown that momentary increases in stress and negative affect precede binge-eating and purging in BN (24, 25) and AN (26), it is unknown if IC mediates this association. Acute stress has increased preference for palatable foods among dieters, which co-occurred with augmented fronto-limbic-striatal functional connectivity and reduced connectivity between the ventromedial prefrontal cortex (vmPFC) and dlPFC (27). Thus, acute stress may impair goal-directed, prefrontal control and instead evoke habitual responding to food. Indeed, a pilot study of BN reported stress-induced decreases in bilateral precuneus, ACC and dlPFC responses to palatable food cues, which moderated the association between stress and binge-eating in daily life (28).

Taken together, there is some evidence of fronto-striatal deficits in BN during self-regulatory tasks; however, the precise IC deficits that subserve binge-eating have not been characterized. Moreover, acute, psychological stress may potentiate binge-eating via impaired IC, yet the impact of stress on neural and behavioral correlates of IC is unknown. We therefore conducted the first examination of the effect of acute stress on two key inhibitory modes—proactive and reactive inhibition—in women with AN-BP, BN and unaffected controls. Participants attended a two-day, inpatient study session, in which they completed repeated fMRI scanning under neutral and stressful conditions. Patient groups were expected to have reactive inhibition inefficiencies at baseline, which would be exacerbated by acute stress and related to reduced inferior frontal and striatal activity. We hypothesized baseline proactive inhibition to be reduced in BN but augmented in AN-BP compared to controls, aligning with restrictive AN (AN-R; (29)). However, we anticipated that both groups would show stress-induced proactive inhibition impairments and correspondingly altered frontoparietal activity. Finally, we conducted exploratory analyses to relate neural and behavioral correlates of inhibitory control to laboratory-based eating behavior.

## 2. Methods and Materials

### 2.1 Participants

We recruited eighty-five women (M±SD = 23.96±3.98y) through posted advertisements, the B-eat charity and an adult ED service in Cambridgeshire. Eligible volunteers were aged 18 to 40 years, English-speaking, had normal or corrected-to-normal vision and, for patient groups, met DSM-*5* diagnostic criteria for either AN-BP or BN. Healthy controls with a lifetime psychiatric disorder were ineligible. Patient volunteers with binge-eating disorder, neurodevelopmental disorders, lifetime serious mental illness, and/or substance or alcohol use disorders (SUDs) in the past 6 months were excluded. Full exclusion criteria are described in the Supplementary Material. The study was approved by the Cambridge East Research Ethics Committee (Ref. 17/EE/0304), and all participants provided signed, informed consent. All procedures were performed in accordance with local regulations.

Participants were matched on age, IQ and, for BN and HC groups, BMI (t(61)=0.19, p=.85; Table 1). Moreover, AN-BP and BN groups reported comparable rates of binge-eating and purging, comorbid psychopathology and medication use (Table S1). Women with AN-BP had a greater lifetime incidence of AN-R (p=.015) while excessive exercise episodes were more frequent in BN (p=.04).

**Table 1.** Clinical and demographic information

| Characteristic | AN (n = 22) | BN (n = 33) | HC (n = 30) | Analysis |  |
| --- | --- | --- | --- | --- | --- |
| | M (SD) | M (SD) | M (SD) | $\chi^2$(df), F(df),<br>W, t(df) | p |
| Age (y) | 24.6 (4.7) | 23.6 (3.9) | 23.9 (3.5) | $\chi^2(2) = 0.8$ | .69 |
| BMI (kg/m<sup>2</sup>) | 16.4 (1.4) | 22.0 (2.4) | 21.9 (2.1) | $\chi^2(2) = 48.4$ | <.001 |
| NART IQ score | 116 (5) | 114 (5) | 114 (5) | $\chi^2(2) = 3.2$ | .21 |
| BDI-II | 35.3 (12.0) | 32.7 (10.5) | 2.4 (2.8) | $\chi^2(2) = 57.7$ | <.001 |
| TAI | 63.1 (10.4) | 62.8 (7.3) | 33.0 (6.9) | $F(2) = 151.1$ | <.001 |
| BIS-11 | 65.9 (12.9) | 67.8 (10.5) | 57.9 (5.7) | $F(2) = 8.6$ | <.001 |
| *Attentional* | 20.4 (4.0) | 19.4 (3.7) | 14.8 (2.6) | $F(2) = 20.5$ | <.001 |
| *Motor* | 21.36 (4.8) | 22.6 (4.4) | 20.8 (2.3) | $F(2) = 1.7$ | .19 |
| *Nonplanning* | 24.2 (6.5) | 25.8 (5.6) | 22.17 (3.2) | $F(2) = 3.8$ | .03 |
| EDE-Q | 4.4 (0.8) | 4.6 (0.8) | 0.2 (0.2) | $\chi^2(2) = 58.0$ | <.001 |
| EDE ratings |  |  |  |  |  |
| *OBEs* | 38.1 (47.9) | 23.0 (29.1) | - | $W = 317.5$ | .43 |
| *SBEs* | 9.5 (12.8) | 6.6 (6.2) | - | $W = 341.5$ | .93 |
| *Vomiting episodes* | 43.5 (51.6) | 24.2 (31.0) | - | $W = 304.0$ | .31 |
| *Laxative episodes* | 1.1 (3.4) | 2.0 (3.9) | - | $W = 421.5$ | .18 |
| *Exercise episodes* | 7.4 (13.6) | 10.9 (9.4) | - | $W = 478.5$ | .04 |
| Age of onset (y) | 15.6 (2.4) | 16.2 (3.1) | - | $t(51.8) = -0.8$ | .42 |
| Illness duration (y) | 9.0 (5.8) | 7.4 (4.0) | - | $t(34.4) = 1.1$ | .27 |
| | N (%) | N (%) | N (%) | $\chi^2$(df) | p |
| Comorbid anxiety | 3 (13.6) | 3 (9.1) | - | $\chi^2(1) = 0.3$ | .69 |
| Comorbid MDE | 15 (68.2) | 16 (48.5) | - | $\chi^2(1) = 2.1$ | .15 |
| Comorbid personality | 2 (9.1) | 5 (15.2) | - | $\chi^2(1) = 0.4$ | .69 |
| Any current treatment | 13 (59.0) | 15 (45.5) | - | $\chi^2(1) = 1.0$ | .32 |
| *Psychotherapy* | 9 (40.9) | 9 (27.3) | - | $\chi^2(1) = 1.1$ | .29 |
| *Medication* | 10 (45.5) | 10 (30.3) | - | $\chi^2(1) = 1.3$ | .25 |
| Prior AN-R | 14 (63.6) | 10 (30.3) | - | $\chi^2(1) = 6.0$ | .01 |

**Note:** BMI = body mass index, NART = National Adult Reading Test, BDI-II = Beck Depression Inventory-II, TAI = Trait Anxiety Inventory, EDE-Q = Eating Disorder Examination Questionnaire, EDE = Eating Disorder Examination, OBE = objective binge-eating episode, SBE = subjective binge-eating episode, MDE = major depressive episode, AN-R = anorexia nervosa restrictive subtype. EDE ratings reflect

### 2.2 Study design

Participants underwent the same study procedure as described previously (30). Briefly, eligible volunteers completed an initial screening session, where they provided informed consent prior to blood sampling and administration of the Eating Disorder Examination (31) and Structured Clinical Interview for DSM-*5* (32) (see Supplementary Material). Fifteen volunteers were excluded following the screening, leaving 85 women (n=22 AN-BP, n=33 BN, n=30 HC) for the two-day, overnight study session. Study sessions began at either 08.00 or 09.00h, and participants’ height and weight were measured prior to a standardized breakfast. Participants then completed a questionnaire and cognitive task battery, and they were offered a mid-morning snack before a 6-hour fast. A cannula was placed approximately 1 hour prior to MRI scanning on Day 1, and blood samples for cortisol and gut hormones were acquired at fixed timepoints (30). Participants began MRI scanning between 13.30 and 14.30h to control for diurnal fluctuations in cortisol. While scanning, participants performed the stop-signal anticipation task (SSAT; (33)) twice, immediately pre- and post-manipulation, and manipulation order (stress vs. neutral) was counterbalanced across participants. Then, participants had an unsupervised ad libitum meal, and those who did not meet their estimated energy requirements were offered an evening snack. The study protocol was identical on Day 2, and participants were discharged following the meal.

### 2.3 Stop-signal anticipation task

The SSAT measures both proactive (i.e., anticipation of stopping) and reactive (outright stopping) inhibition. The task and procedure have been described previously (33), and an overview is included in Figure 1 and the Supplementary Material. Code may be retrieved from: https://github.com/bramzandbelt/SSAT.

**Figure 1.**
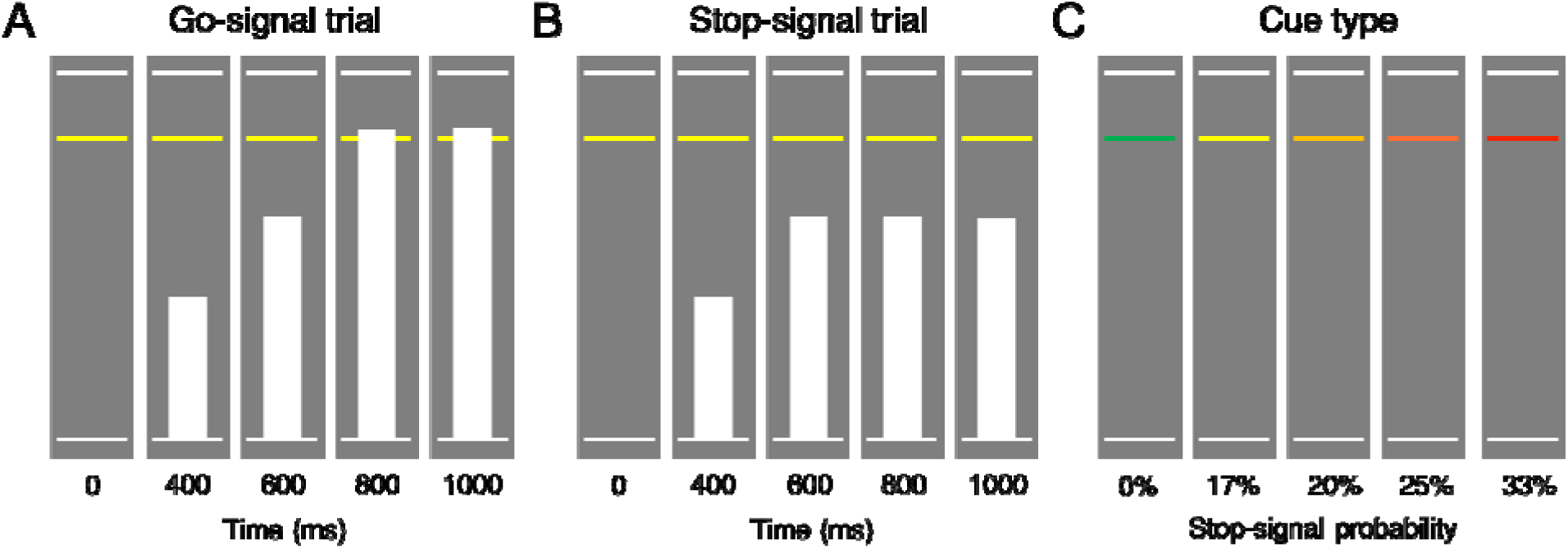
Schematic of stop-signal anticipation task. Schematic of SSAT trial types adapted from Zandbelt & Vink (33). **A)** On go-signal trials, participants were instructed respond when a moving bar reached the middle line. The target response time was 800ms on each 1000ms trial (1000ms inter-trial interval). **B)** A minority of trials (25%) were stop-signal trials, where the moving bar stopped automatically before reaching the middle line. Participants were instructed to withhold their response in the event of a stop-signal. **C)** To index proactive inhibition, the probability of a stop-signal occurring on a given trial ranged from 0 – 33%, as indicated by colored cues. Participants were told that stop-signals would never occur on “green” (baseline) trials, but the likelihood of a stop-signal occurring increased across yellow to red trials.

### 2.4 Stress induction

On each day, participants completed either an acute, psychological stress induction or a control task (i.e., neutral condition) to enable within-subject assessment of stress responses. Details of the task structure have been described elsewhere (https://github.com/mwestwater/STRIvE-ED; (30)). Participants completed 48 multiple-choice, mental math problems of varying difficulty in each condition; however, in the stress induction, participants were motivated to respond accurately whereas performance was not evaluated during the control task. Moreover, incorrect responses elicited negative feedback (e.g., *“Your performance is below average.”*) in the stress task, and uncontrollability, a central aspect of psychological stress, was engendered through the delivery of mild electrical stimulation to the abdomen at variable frequencies and intensities. Importantly, subjective ratings of stimulation intensity, unpleasantness and pain did not differ significantly across groups, indicating that abdominal stimulation was suitable for ED participants (see Supplementary Material). Subjective stress ratings were collected immediately pre- and post-induction.

### 2.5 Image acquisition

MR images were acquired on a 3T Siemens Skyra^Fit^ scanner (Erlangen, Germany) fitted with a 32-channel, GRAPPA parallel-imaging head coil. MR sequences are described in the Supplementary Material. One participant was excluded for an incidental finding of white matter abnormalities.

### 2.6 Data analysis – SSAT performance

We assessed proactive inhibition by examining the effect of stop-signal probability on response time (RT), where participants tend to slow responding as the likelihood of having to stop increases (33–36). Impaired proactive inhibition would be evident in a failure to increase RT when stop-signal probability increases, as this would suggest weaker anticipation of stopping. Reactive inhibition was indexed as stop-signal reaction time (SSRT), which represents the latency of the inhibition process. SSRT was computed using the integration method (37) across all stop-signal probability levels. Slower SSRTs would reflect greater latency of the inhibitory process and therefore impaired reactive inhibition.

Behavioral data were analyzed in R (38). Aligning with previous reports (33, 39), go-signal RTs that were more than 1.5 times the interquartile range below the 25^th^ percentile or above the 75^th^ percentile of the RT distribution at each probability level, as well as on failed stop-signal trials, were defined as outliers. To minimize positive skew, a rank based inverse normal transformation was applied to RTs (R package *RNOmni* (40)) prior to analysis. Analyses of proactive inhibition (trial RT) and reactive inhibition (SSRT) were conducted using the linear mixed-effects modelling (LMM) R package *nlme* (41), where fixed effects of group, condition and time were included in both models, with random intercepts for within-subject variables nested within the subject’s random effect. Additionally, fixed and random effects for probability level (linear and quadratic terms) were included in the proactive inhibition LMM. Normality of the model residuals was determined by visual inspection of quantile-quantile plots.

### 2.7 Data analysis – fMRI

Image data were pre-processed and analyzed using FreeSurfer (v 6.0; (42, 43)) and AFNI software (44). Pre-processing steps were completed with the afni_proc.py python script with 6mm spatial smoothing (see Supplementary Material for details). Statistical analysis followed a two-level procedure, where successful stop-signal trials, failed stop-signal trials, go-signal trials with non-0% stop-signal probability were modelled as regressors of interest in the first-level general linear models. In line with previous work (33, 39), we included two amplitude modulators, RT and stop-signal probability level, for go-signal trials. AFNI models one regressor for the constant magnitude of the blood oxygenation-dependent (BOLD) response and separate regressors for each amplitude per time point unlike other packages that partition the variance of regressors sequentially. However, as RT (a measure of the tendency to withhold a response) and stop-signal probability contrasts may provide complementary information, both were used as measures of proactive inhibition. In addition, incorrect go-signal trials and rest blocks were included as nuisance regressors; go-signal trials with a stop-signal probability of 0% were not modelled, thus constituting an implicit baseline. Regressors were created by convolving gamma functions coding for response onset (or target RT for successful stop-signal trials) with a canonical hemodynamic response function. Within each subject run, we computed four contrast images: 1) the parametric effect of RT on go-signal activation (proactive inhibition), 2) the parametric effect of stop-signal probability on go-signal activation (proactive inhibition), 3) successful stop versus failed stop-signal trials (reactive inhibition) and 4) successful stop versus go-signal trials with 0% stop-signal probability (reactive inhibition). We generated two contrasts for reactive inhibition as there is no consensus on which contrast is most appropriate when investigating this inhibitory mode. Beta estimates were determined using restricted maximum likelihood estimation.

We conducted two group analyses for each contrast. First, we examined associations between diagnostic group, condition (stress vs. neutral), time (pre vs. post) and their interaction and the BOLD response in seven predefined regions of interest (ROIs) previously implicated in proactive inhibition and reactive inhibition (Figure 2A & Supplementary Material). For each ROI, main and interaction effects were tested in a LMM, and random intercepts for condition and time were included within the random effect of the individual. As seven ROIs were tested per contrast, our alpha threshold was reduced to p=.05/7=.007. Next, we examined whether a three-way interaction between group, time and condition related to differences in whole-brain activation. Whole brain analyses were completed using the linear-mixed effects modelling AFNI program, *3dLME* (45), where general linear tests were implemented to test a priori contrasts of interest, including AN>HC, BN>HC, stress>neutral, post>pre and the three-way interaction (e.g., AN>HC*stress>neutral*post>pre). For completeness, both F- and Z-statistics are reported for each effect. Resulting group-level statistical maps were tested for significance using cluster-level inference (cluster-defining threshold of p<.001, k=18.8, cluster probability of p<.05, family wise error-corrected). Updated versions of *3dFWHMx* and *3dClustSim* were used to correct for multiple comparisons, as these programs incorporate a mixed autocorrelation function to model non-Gaussian noise structure and reduce false-positive rates (46, 47). For visualization, the mean percent signal change was extracted from significant whole-brain clusters using *3Dmaskave.*

**Figure 2.**
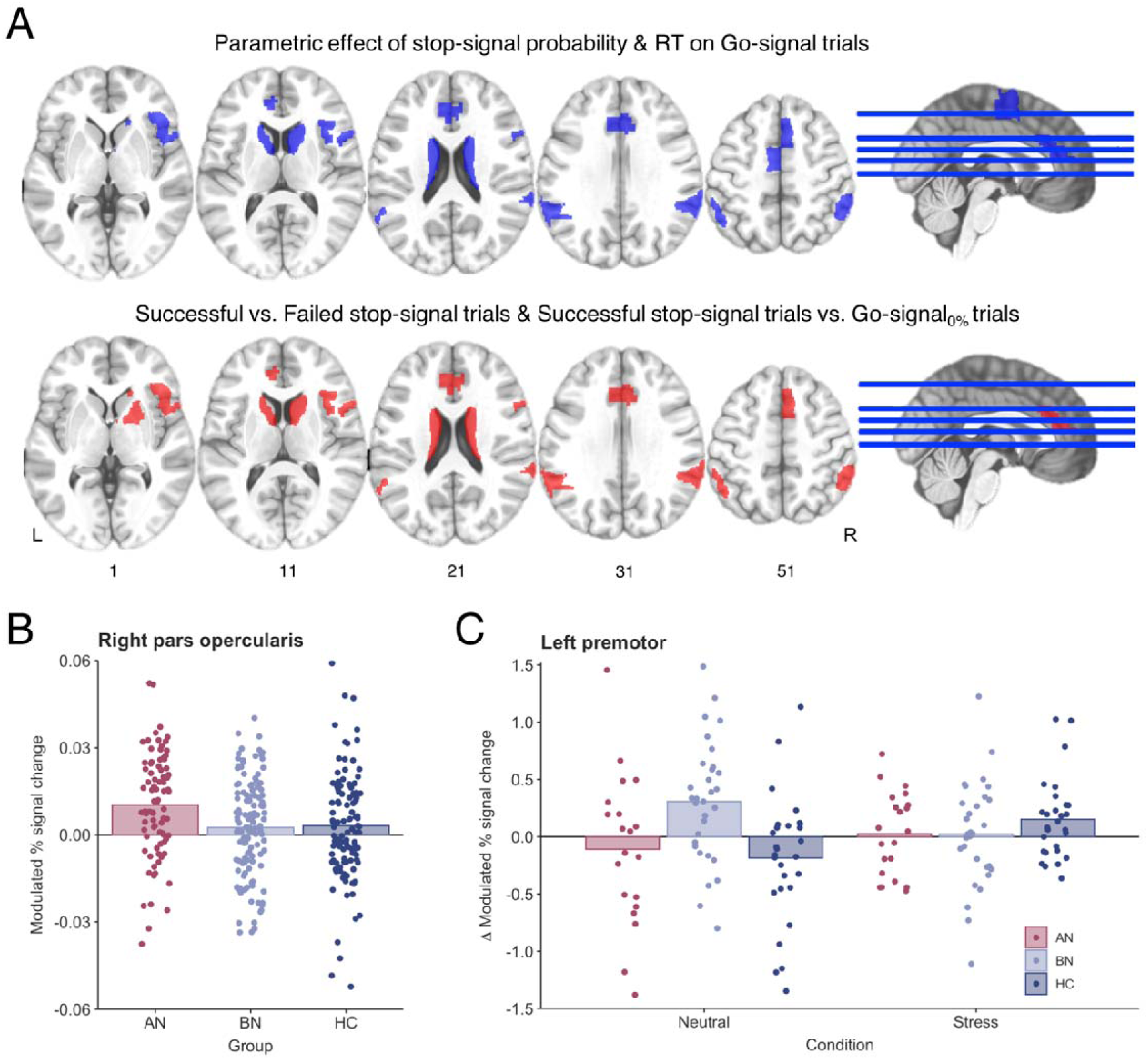
Region of interest analyses identify altered inferior frontal and premotor activity during proactive inhibition in anorexia and bulimia nervosa. **A)** ROI analyses were conducted in seven regions that have previously been associated with proactive (blue) and reactive (red) inhibition (22, 39). The following ROIs were used in analysis of both proactive and reactive inhibition: right opercular inferior frontal gyrus, right ventral inferior frontal gyrus, bilateral caudate, bilateral pregenual anterior cingulate cortex, right pre-supplementary motor cortex and bilateral superior parietal cortices. Analysis of proactive inhibition also included the left premotor cortex, and the right putamen was included in reactive inhibition analysis. ROIs are displayed in neurological orientation (L=left). **B)** The parametric effect of stop-signal probability was related to increased right inferior frontal gyrus (pars opercularis) activity in AN-BP relative to controls (p=.005). **C)** A three-way interaction indicated that the parametric effect of reaction time was related to increased left premotor activity in BN compared to controls in following the control task.

### 2.7 Exploratory analysis of inhibitory control and ad libitum consumption

We used LMMs to test whether SSRT, Barratt Impulsiveness scores (BIS-11; 48) or brain regions implicated in the fMRI analyses explained variance in subsequent food intake (see Supplementary Material). Exploratory results were considered statistically significant at p=.05.

## 3. Results

### 3.1 Behavioral

#### 3.1.1 Manipulation check

As previously reported, both subjective stress and negative affect were significantly increased following the stress induction relative to the control condition (Figure S1). Moreover, a group-by-condition interaction identified stress-induced plasma cortisol decreases in BN, but not AN-BP, compared to controls (30).

#### 3.1.2 Reduced proactive inhibition in bulimia nervosa

We anticipated that proactive inhibition would be impaired in BN and augmented in AN-BP while stress-induced impairments would be observed in both groups. RT increased with greater stop-signal probability (β=0.01, t(1019)=13.08, p<.0001); however, this effect was nonlinear, as a significant quadratic probability term suggested that RT slowing plateaued with increasing stop-signal probability (β=−5.07, t(57919)=−5.28, p<.0001). RT on non-0% go-signal trials was significantly decreased post-manipulation (i.e., at time 2; β=−0.14, t(169)=−8.49, p<.0001), which is consistent with the expected practice effects on each day. Moreover, a significant group-by-probability interaction indicated poorer proactive inhibition in the BN group relative to controls (β=−6.54, t(1012)=−2.97, p=.003; Figure 3A), where women with BN demonstrated a smaller increase in RT relative to increasing stop-signal probability. The addition of higher-order interaction terms did not significantly improve model fit (χ^2^(13)=16.11, p=.19), indicating that proactive inhibition was not significantly affected by acute stress. RT on 0% stop-signal probability trials did not differ between AN (p=.37) or BN (p=.96) and control participants, indicating equivalent performance on the baseline response task (Table S2).

**Figure 3.**
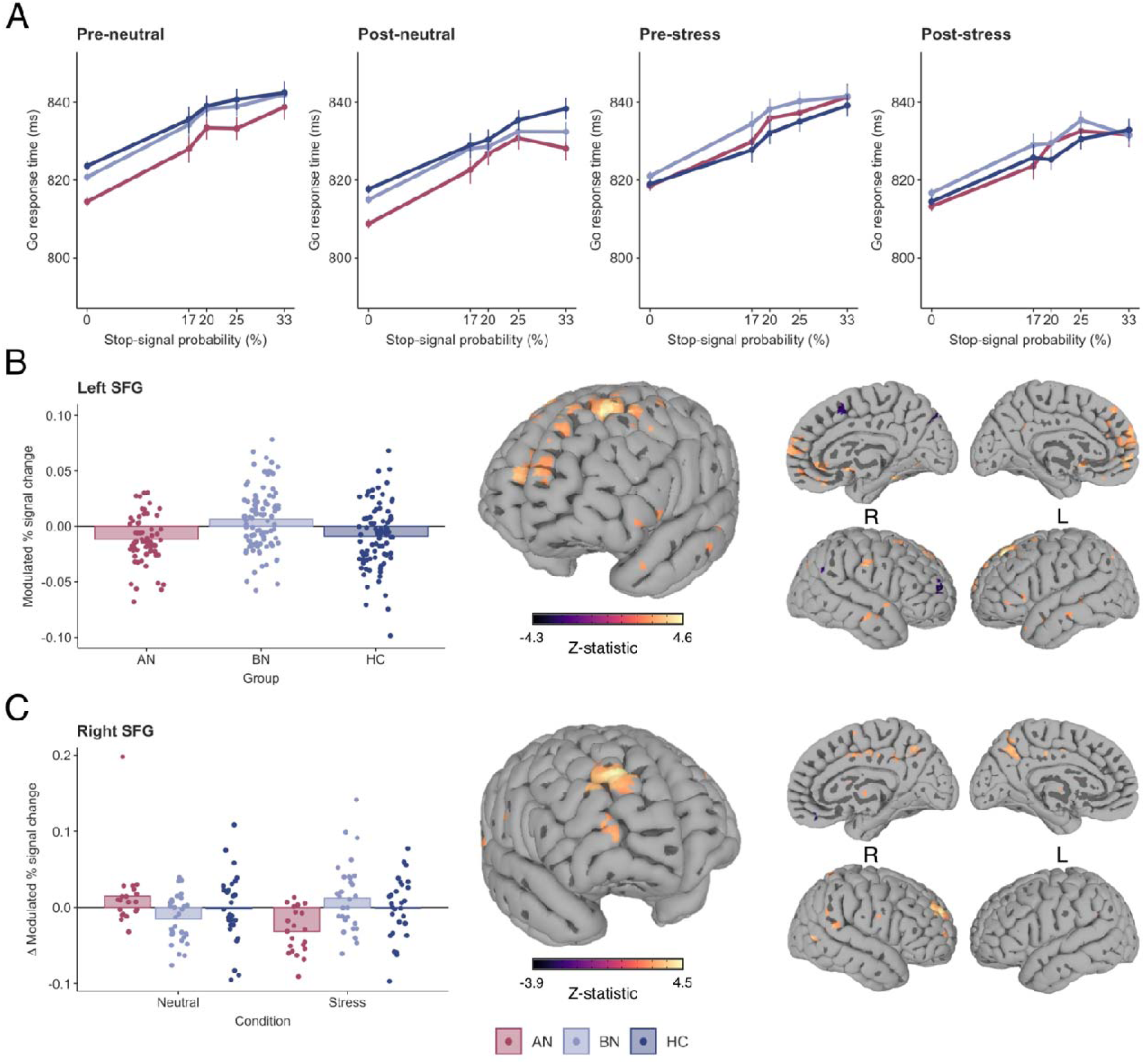
Impaired proactive inhibition in bulimia nervosa is associated with increased superior frontal gyrus activity. **A)** Reaction time increased as a function of stop-signal probability in all groups; however, a significant group-by-probability interaction showed that women with BN did not slow to the same degree as controls in response to increasing stop-signal probability (p=.003). This impairment in proactive inhibition was associated with greater activity in **B)** the left superior frontal gyrus (k = 25 voxels, Z = 4.58, MNI_X,Y,Z_ = −23, 33, 54, cluster defining threshold = p<.001, FWE-corrected cluster probability = p<.05) in BN relative to controls. **C)** A three-way interaction was related to stress-induced increases in the right superior frontal gyrus in BN relative to AN-BP (k = 34 voxels, Z = 4.52, MNI_X,Y,Z_ = 22, 54, 36, cluster defining threshold = p<.001, FWE-corrected cluster probability = p<.05). For illustration, whole-brain activation was thresholded at voxel-wise p<.01 (uncorrected). Individual values are overlaid on the mean modulated % signal change by group. Error bars = SEM.

#### 3.1.3 No effect of patient group or stress on reactive inhibition

We predicted that both AN-BP and BN groups would demonstrate impaired reactive inhibition relative to controls following the acute stress induction. The significant main effect of time indicated that SSRT was reduced post-manipulation (β=−3.29, t(166)=−3.23, p=.002). However, all other main and interaction effects on SSRT were nonsignificant (all p’s>.05; Table S2). Data met the assumptions of the race model, as evidenced by faster RTs on failed stop-signal trials compared to go-signal trials where stop-signals could occur (β=−21.5, t(339)=−39.4, p<.0001).

### 3.2 Functional MRI

#### 3.2.1 Proactive inhibition

Examination of the parametric effects of stop-signal probability and RT identified increased neural responses across frontoparietal regions that comprise the proactive inhibition network (Tables S4-5, Figure S2), indicating successful experimental manipulation of proactive inhibition.

##### ROI analyses

Increasing stop-signal probability was associated with greater right inferior frontal gyrus activity in the AN-BP group relative to controls (β=0.007, t(81)=2.91, p=.005; Figure 2B). IFG activity decreased post-manipulation (i.e., at time 2) across all groups (β=−0.006, t(156)=−3.20, p=.002). In addition, the parametric effect of RT on left premotor cortex activity was related to a three-way interaction, where the BOLD response decreased in BN relative to controls following the stress induction (β=−0.62, t(151)=−3.48, p<.001; Figure 2C).

##### Whole-brain analyses

Increasing RT was related to reduced left supplementary motor area (SMA) activity post-manipulation (Table 2). Moreover, the effect of stop-signal probability was significantly affected by time, where activity across the proactive inhibition network generally decreased post-manipulation (Table 2). In line with behavioral findings, the effect of stop-signal probability also differed significantly by group, where the parameter was related to increased activity in the left superior frontal gyrus (SFG) in BN relative to controls (k=25 voxels, Z=4.58; Figure 3B & Table 2). A significant three-way interaction was associated with right SFG activity (k=19 voxels, F(2,231)=10.77). As this effect was not captured by our a priori contrasts, we conducted an additional general linear test, which indicated augmented SFG activity in BN relative to AN-BP following stress (k=34 voxels, Z=4.52; Figure 3C & Table 2).

**Table 2.**
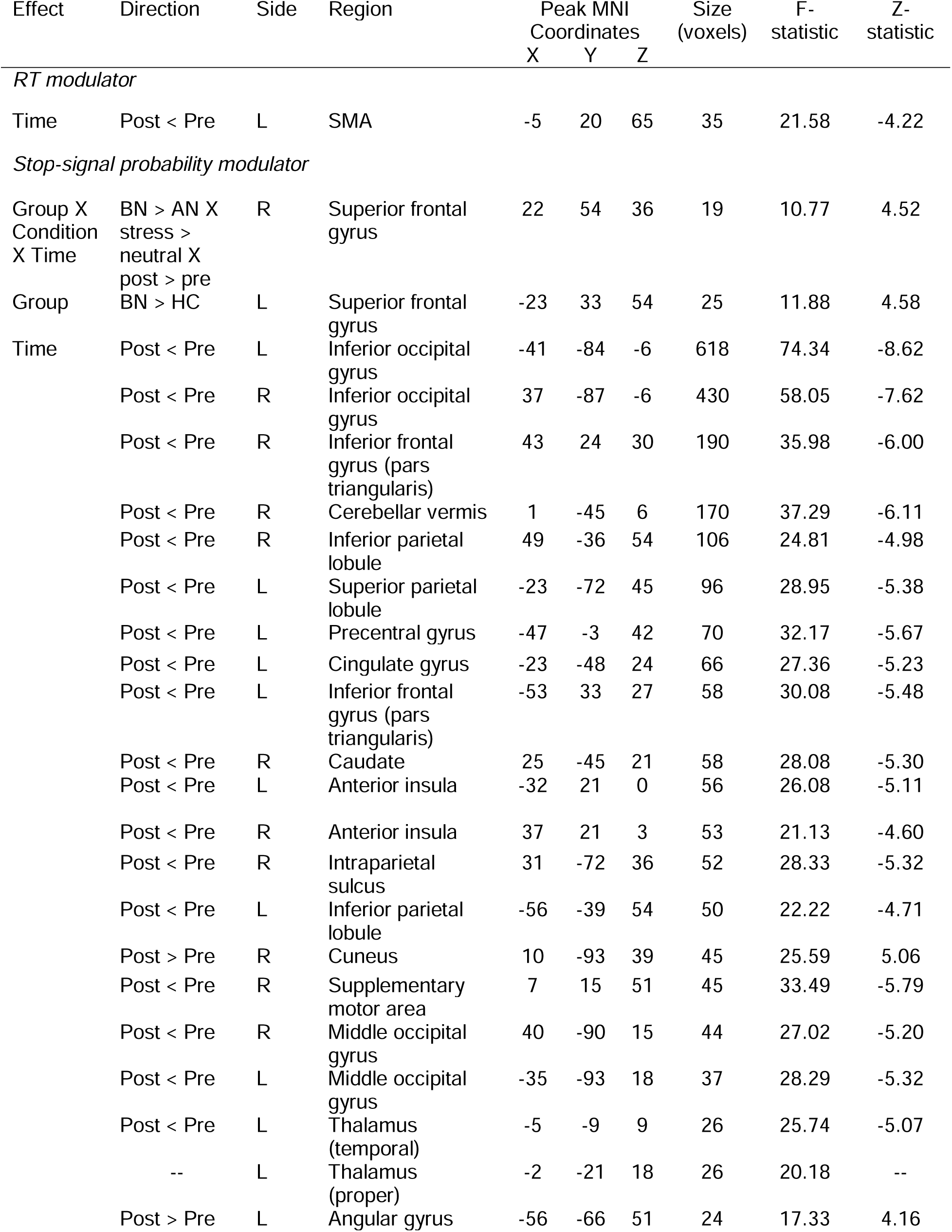
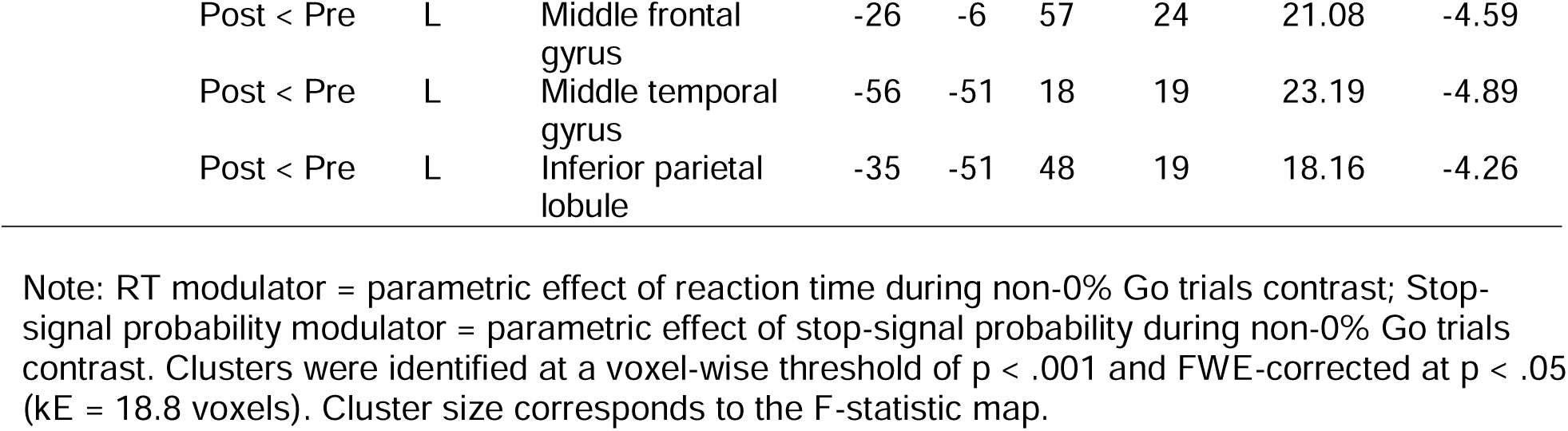
Whole-brain derived fMRI responses during proactive inhibition

#### 3.2.2 Reactive inhibition

Analyses of reactive inhibition (Stop>Go-signal and Stop>FailedStop trials) indicated increased neural responses across the IC network (Tables S6-7, Figure S3) with markedly similar activation patterns across groups.

##### ROI analyses

The main effect of group and all interaction effects were nonsignificant across all ROIs for both reactive inhibition contrasts. Significant main effects of time and condition (i.e., scan day) are reported in the Supplementary Material.

##### Whole-brain analyses

On Stop>Go-signal trials, neural responses were significantly reduced across the IC network post-manipulation (Table S3). Activity in left middle temporal, thalamic, posterior insular, occipital and inferior frontal clusters was reduced post-manipulation during Stop>FailedStop trials. Moreover, left precentral gyrus activity on Stop>FailedStop trials was increased on the stress day relative to the neutral day. Finally, a three-way interaction indicated reduced activity in the right vmPFC during reactive inhibition (Stop>FailedStop trials) in AN-BP relative to controls following stress (k=32 voxels, Z=−4.19; Figure 4 & Table 3).

**Table 3.** Whole-brain derived fMRI responses during reactive inhibition (Successful Stop > Fail Stop)

| Effect | Direction | Side | Region | Peak MNI Coordinates |  |  | Size (voxels) | F-statistic | Z-statistic |
| --- | --- | --- | --- | --- | --- | --- | --- | --- | --- |
|  |  |  |  | X | Y | Z |  |  |  |
| Group X Condition X Time | AN > HC X Stress > Neutral X Post > Pre | R | Medial frontal gyrus | 4 | 45 | -9 | 30 | 12.35 | -4.19 |
| Condition | Stress < Neutral | L | Precentral gyrus | -59 | -3 | 33 | 24 | 16.16 | -4.02 |
| Time | Post < Pre | L | Middle temporal gyrus | -59 | -60 | 18 | 37 | 14.90 | -3.86 |
|  | Post < Pre | L | Thalamus (prefrontal) | -8 | -18 | 12 | 21 | 17.68 | -4.21 |
|  | Post < Pre | L | Posterior insula | -44 | -3 | 0 | 21 | 16.33 | -4.04 |
|  | Post < Pre | L | Superior occipital gyrus | -20 | -78 | 33 | 19 | 15.53 | -3.94 |

Note: Clusters were identified at a voxel-wise threshold of $p < .001$ and FWE-corrected at $p < .05$ ( $kE = 18.8$ voxels). Cluster size corresponds to the F-statistic map.

**Figure 4.**
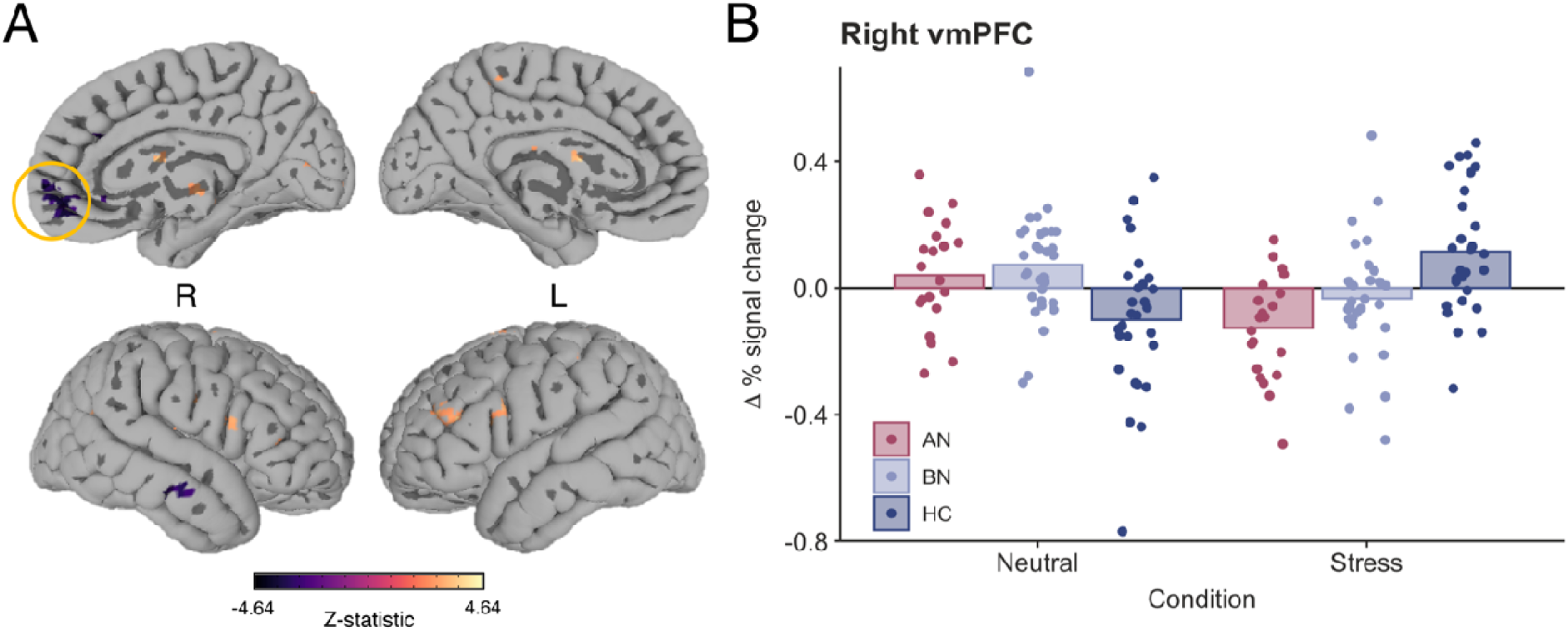
Stress reduces right ventromedial prefrontal cortex activity in anorexia nervosa (binge/purge subtype) during reactive inhibition. **A)** A significant three-way interaction indicated that right vmPFC activity was significantly reduced following acute stress compared to the neutral condition in AN-BP relative to controls (k = 32 voxels, Z = −4.19, MNI_X,Y,Z_ = 4, 45, −9, cluster defining threshold = p < .001, FWE corrected cluster probability = p < .05). Whole-brain activation was thresholded at voxel-wise p < .01 (uncorrected) for illustration. **B)** Change in average percent signal change for the vmPFC cluster from pre- to post-induction across conditions. Individual values are overlaid on the mean change in percent signal change (post – pre) by group.

#### 3.2.3 Associations with food intake

We previously reported that AN-BP and BN groups consumed less in the buffet than controls, and intake was unaffected by stress (30). Here, exploratory analyses linked left SFG responses during proactive inhibition to increased intake (Z-scored; β=3.59, t(70)=2.36, p=.02), vmPFC responses during reactive inhibition were negatively related to consumption (β=−0.81, t(70)=−2.84, p=.006; Figure S4). The effects of SSRT, trait impulsivity and all interaction terms were nonsignificant (all p’s>.05).

## 3. Discussion

As failed self-regulation in response to stressors has gained traction as a putative mechanism of binge-eating, it has become increasingly important to characterize the precise self-regulatory deficits associated with binge-eating disorders. We assessed the impact of induced stress on IC in women with AN-BP, BN and matched controls, reporting three key findings. First, women with BN, but not AN-BP, had impaired proactive inhibition, yet both groups demonstrated increased prefrontal responses during the anticipation of stopping compared to controls. Second, we found stress-induced changes in the neural correlates of proactive and reactive inhibition, with notable differences across diagnostic groups. Third, AN-BP and BN groups had intact reactive inhibition, and neither proactive nor reactive inhibition performance was affected by acute stress.

We report novel evidence of impaired proactive inhibition in BN relative to controls, which co-occurred with increased activity in the left dorsolateral SFG. Increased left SFG activity and concurrent performance deficits could reflect inefficient recruitment of other regions within the IC network, namely inferior and middle frontal gyri, which share reciprocal connections with the SFG (49). Inefficient or compensatory responses may also explain increased right IFG responses in AN-BP during intact proactive inhibition. Alternatively, given the role of the pars opercularis in ‘braking’ motor responses (50, 51), increased activity could reflect improved proactive adjusting in AN-BP on the neural level, complementing previous behavioral reports in AN-R (29). Exploratory analyses found that left SFG responses predicted increased post-scan calorie intake, lending additional support to the notion of inefficiencies across the proactive inhibitory network that may relate to disordered eating behavior.

Acute, psychological stress altered right SFG and left premotor cortex responses during proactive inhibition, as well as right vmPFC activity during outright stopping, differently between groups. Specifically, stress augmented right SFG responses to increasing stop-signal probability in BN relative to AN-BP. In BN, these stress-induced increases in SFG responses perhaps compensated for concomitant decreases in premotor activity during RT slowing, thus preserving task performance. Indeed, increased prefrontal activity has been reported in healthy adults following pain stress, where activation was presumed to support working memory performance (52).

One explanation for augmented vmPFC responses in controls relative to AN-BP after stress could be stress-induced alterations in inter-regional modulation (53). The vmPFC is the primary cortical target of limbic projections (54), and stress-induced increases in activity may provide top-down modulation of amygdala reactivity and negative emotions. While not typically associated with inhibitory control, augmented vmPFC activity during reactive inhibition has been reported following methylphenidate administration (55) and neuromodulation of the pre-SMA (56). These findings, together with our observations following acute stress, could implicate norepinephrine signaling in altered vmPFC activation, but further research is needed. Our finding of a negative relationship between vmPFC responses to reactive stopping and post-scan calorie consumption suggests that vmPFC activation during IC may be important for dietary control.

Stress-induced reductions in prefrontal responses during both proactive and reactive inhibition in AN-BP could reflect the consequences of prolonged, extreme stress, namely significantly low weight, which engenders various cognitive and neuroendocrine perturbations (57, 58). Interestingly, preclinical research has identified disrupted dopaminergic signaling following severe stress (59, 60); however, the effect of stress on dopaminergic projections to prefrontal cortex remains understudied. The dearth of research in this area discourages a premature interpretation of our stress induction effects in AN-BP. Instead, findings of task-specific, stress-induced reductions in prefrontal responses in AN-BP may inform future investigations into neurocognitive alterations associated with prolonged and increasing stress.

Contrary to our hypotheses, reactive inhibition, indexed as SSRT, was unaffected by diagnostic group or stress, and it was unrelated to free-choice consumption. As we have reviewed, findings of impaired self-regulatory performance in BN and AN-BP are inconsistent (10, 12), and our results suggest that the subjective ‘loss of control’ that characterizes binge-eating episodes does not relate to deficits in one’s capacity for action cancellation. While often considered a valid and translational measure of IC, our findings, and a recent mega-analysis of polysubstance use, question the clinical utility of SSRT. Indeed, the latter found that increased SSRT was not significantly related to various SUDs, including alcohol and cocaine use disorders (61). As stress-induced deficits in the ability to delay food reward were found in non-clinical samples (62), future research should assess state changes in decision-making as a potential mechanism of loss-of-control eating in clinical groups.

Although our design had notable strengths, several limitations should be considered. First, we recruited a representative sample of women with EDs, and as expected, the majority suffered with comorbid psychopathology and many used medication. These characteristics may, however, improve the generalizability of our findings as comorbidity and medication use are the norm rather than the exception amongst individuals with EDs (3, 63). Moreover, of those using medication, most were prescribed either selective serotonin reuptake inhibitors or serotonin-norepinephrine reuptake inhibitors with high affinity for 5-HT, and 5-HT modulation has been shown to have no effect on response inhibition (64). Second, disorder-salient stimuli (e.g., food), which may accentuate or reveal self-regulatory deficits (65), were not used, and future study should examine the impact of stress on performance in these contexts. Third, the conditions under which stress was induced (i.e., in an MR scanner) and eating behavior was assessed differed from those in daily life.

Our findings counsel against a simplistic, stress-induced failure of regulation as an overall explanation for binge-eating in AN-BP and BN, underscoring the need for alternative models of these illnesses. Moreover, dissociations across diagnostic groups suggest that models of binge-eating based on BN may not apply to AN-BP. Given the complex metabolic and psychological disturbances associated with these disorders, future efforts to identify the neurocognitive mechanisms of binge-eating should consider the roles of interacting peripheral physiological processes.

## Acknowledgements

The authors wish to thank the participants for their time and dedication to this project; the nursing staff and radiographers for assisting the study protocol; and Dr. Bram Zandbelt for sharing the stop-signal anticipation task code. They also thank the Beat eating disorder charity for advertising the study. Funding was provided by the Bernard Wolfe Health Neuroscience Fund to PCF and HZ and a Wellcome Trust Investigator Award to PCF (Reference No. 206368/Z/17/Z). MLW was supported through the NIH-Oxford-Cambridge Scholars Program and a Cambridge Trust fellowship. FM was supported by research grants from Versus Arthritis, the Experimental Psychological Society and a Career Development Award from the Medical Research Council (MR/T010614/1). AG, CG and ME were supported by the Intramural Research Program of the NIMH (Ref. ZIAMH002798). The Wellcome Trust/NIHR Clinical and Translational Research Facilities and the Wolfson Brain Imaging Centre provided equipment and support staff for the study.

