## Supplementary Material for "Prefrontal responses during proactive and reactive inhibition are differentially impacted by stress in anorexia and bulimia nervosa"

1. **Supplementary Methods**

***1.1 Screening protocol and exclusion criteria***

As previously described (1), potential volunteers completed a telephone screening and self-report questionnaire of psychopathology symptoms (2) prior to attending an outpatient screening session at Addenbrooke’s hospital, Cambridge, UK. At the screening, participants provided written, informed consent, and a fasting blood sample was collected to assess full blood count and thyroid hormones. Participants’ height, weight and body composition were measured prior to a clinical assessment, where the Eating Disorder Examination (v16; 3) and the Structured Clinical Interview for DSM-*5* (4) were administered to determine ED diagnoses and comorbid psychopathology, respectively. Participants also completed the National Adult Reading Test (5) to determine their estimated IQ, and the Fagerström Test for Nicotine Dependence (6) was administered to assess nicotine dependence. To reduced participant burden, patient participants who lived outside of Cambridgeshire (n=12) completed the screening session remotely. Participants who underwent remote screening completed all blood sampling and anthropometric measurements during the overnight study session.

For all groups, exclusion criteria included: left handedness, estimated IQ<80, body mass index (BMI)>29.9 kg/m^2^, MRI contraindications (e.g., pregnancy, some metallic implants), metabolic, neurological or cardiovascular diseases (e.g., anemia), lactation, bariatric surgery, and high nicotine dependence, as per the Fagerström Test for Nicotine Dependence.

***1.2 Stop-signal anticipation task***

The stop-signal anticipation task (SSAT) was presented using Presentation software (v. 20; Neurobehavioral Systems). A background of three horizontal lines was present throughout the task. On each trial, a bar moved at a constant speed from the bottom line, reaching the top line in 1000ms. The main task (i.e., go-signal trials) involved stopping the moving bar as it reached the middle line with one’s right index finger, yielding a target response time of 800ms. On stop-signal trials, the moving bar stopped automatically before reaching the middle line, signaling that a motor response had to be suppressed. The color of the middle line indicated the probability of a stop-signal occurring on a given trial, where green = 0%, yellow = 17%, amber = 20%, orange = 25% and red = 33%.

The initial stop-signal onset time was set to 500ms (i.e., 300ms before the target response time) for each stop-signal probability level. Throughout the task, the stop-signal onset time was adjusted using a staircase procedure (with steps of 25ms) depending on stopping accuracy. These adjustments were made separately across each stop-signal probability level, ensuring roughly equal numbers of successful and failed stop-signal trials.

Trials were presented in either baseline or experimental blocks that were comprised of 12 to 15 trials each. The inter-stimulus interval was 1000ms. During baseline blocks, participants responded to trials in which the stop-signal probability was 0%, as indicated by the green stop-signal probability cue. Experimental blocks were comprised of go-signal trials with a stop-signal probability >0% (i.e., non-green cues) and stop-signal trials (also non-green cues). Stop-signal trials occurred pseudo-randomly throughout experimental blocks, and stop-signal probability level varied across trials. Distinct trial orders were used for pre- and post-induction runs to account for practice effects within each day, where the trial orders were the same across all participants and scan sessions. Simulations to determine the optimal trial order indicated that correlations between the different model regressors were sufficiently weak to generate parameter estimates.

In total, the SSAT included 474 trials: 234 go-signal trials with a stop-signal probability of 0%, 180 go-signal trials with a stop-signal probability >0% and 60 stop-signal trials. In other words, the proportion of stop-signal trials was 33.3%. Two, 24s rest blocks were presented after one-third and two-thirds of the trials had elapsed. The task duration was 16min 36s. Participants completed a behavioral practice session prior to fMRI scanning on Day 1, in which they were trained on the Go and Stop tasks. Participants were notified that it was equally important to stop the moving bar at the target and withhold their response in the presence of a stop-signal. We informed participants that stop-signals would never occur on trials with green cues, and the likelihood of a stop-signal occurring was lowest on ‘yellow’ cue trials and highest on ‘red’ cue trials, increasing as the cue color transitioned to red. On Day 2, participants were reminded of the task instructions prior to scanning.

***1.3 Acute stress induction***

As previously described (1), the acute stress induction and control task required participants to complete 48 multiple-choice math problems of varying difficulty. Task stimuli were presented in MATLAB (v2017b; The Mathworks), using Psychophysics Toolbox (v3; 2), and code may be retrieved from: <https://github.com/mwestwater/STRIvE-ED>. Prior to the stress induction, participants were encouraged to respond accurately, and they were told that only data from participants whose performance met the group average could be used in the study. Additionally, they were informed that ‘physical distractors’ would be delivered to their abdomen, and that they would be watched on a video camera to ensure they paid attention to the task. Conversely, participants were told that their performance would not be evaluated and that they would not be watched during the control task.

On each day, participants completed 25 practice problems of varying difficulty, and they were instructed to try their best to select the correct answer without taking too much time. Stimuli were presented for a maximum of 30s, during which participants had to respond by selecting one of the 3 choices. Feedback (2500ms) was presented either 500ms after the response, or after the 30s period. The next trial was presented following a variable interval (500 – 2500ms, jitter = 100ms).

Both the stress induction and control task had the same trial structure as the practice task; however, for the stress induction, the initial stimulus presentation and response time (30s in the practice task) was set to 10% less than the participant’s average response time during the practice session. Accurate responses on 3 consecutive trials shortened the maximal response window by 10%, ensuring low performance. As the sliding response window reduced the overall task duration, the ITI was set to 6s on every 6^th^ trial to ensure that the task was sufficiently long for the stress induction to be effective. Participants received negative feedback to nonresponses and incorrect responses (e.g., *“Your performance is below average”*), whereas no feedback was provided for correct responses. At the end of the task, participants were informed that their performance did not meet the group average. For the control task, the stimulus presentation and response time was 30s on each trial, and feedback was only provided to indicate correct responses.

Throughout each task, ‘physical distractors’ were delivered to the abdomen in the form of mild electrical stimulation, using a DS7A constant current stimulator (Digitimer, UK). Prior to MRI scanning, the intensity of electrical stimulation was calibrated for each participant to control for individual differences in shock tolerance. Two BIOPAC radio translucent electrodes (EL509) were filled with isotonic paste (GEL101) and placed 1 – 2 inches to the right of the participant’s navel, between dermatomes T10 and T12. During the calibration procedure, participants indicated (1) when the stimulation was detectable but not uncomfortable and (2) the first moment when the stimulation was uncomfortable but not painful, corresponding to pain ratings of 0 – 2 and 5 – 7, respectively (0 = *no pain*, 10 = *very painful*). Each shock pulse lasted 500μs.

For the stress induction, shocks were delivered in 5 – 20 pulse sequences with an inter-pulse interval range of 0.1 – 1s and an inter-train interval range of 0.1 – 3.9s, which were randomly sampled in MATLAB. Intensity was manually adjusted between the participant’s two threshold values. For the control task, stimulation was delivered at predictable intervals and a constant intensity, which corresponded to the participant’s detection threshold. Trains consisting of 5 pulses were delivered at an inter-pulse interval of 0.55s with an inter-train interval of 2s. Shock delivery was not contingent on performance. Participants were asked to verbally indicate if the stimulation became painful at any time during the task, in which case it would be reduced. No participants reported discomfort.

Participants provided subjective ratings of the stimulation immediately following the task. As expected, pain, intensity and unpleasantness were significantly increased following the stress induction relative to the control task (all p’s <.001). Both the main effect of group and a group-by-condition interaction term were nonsignificant, indicating that women with eating disorders did not perceive the stimulation to be more aversive or distressing than unaffected women perceived it to be.

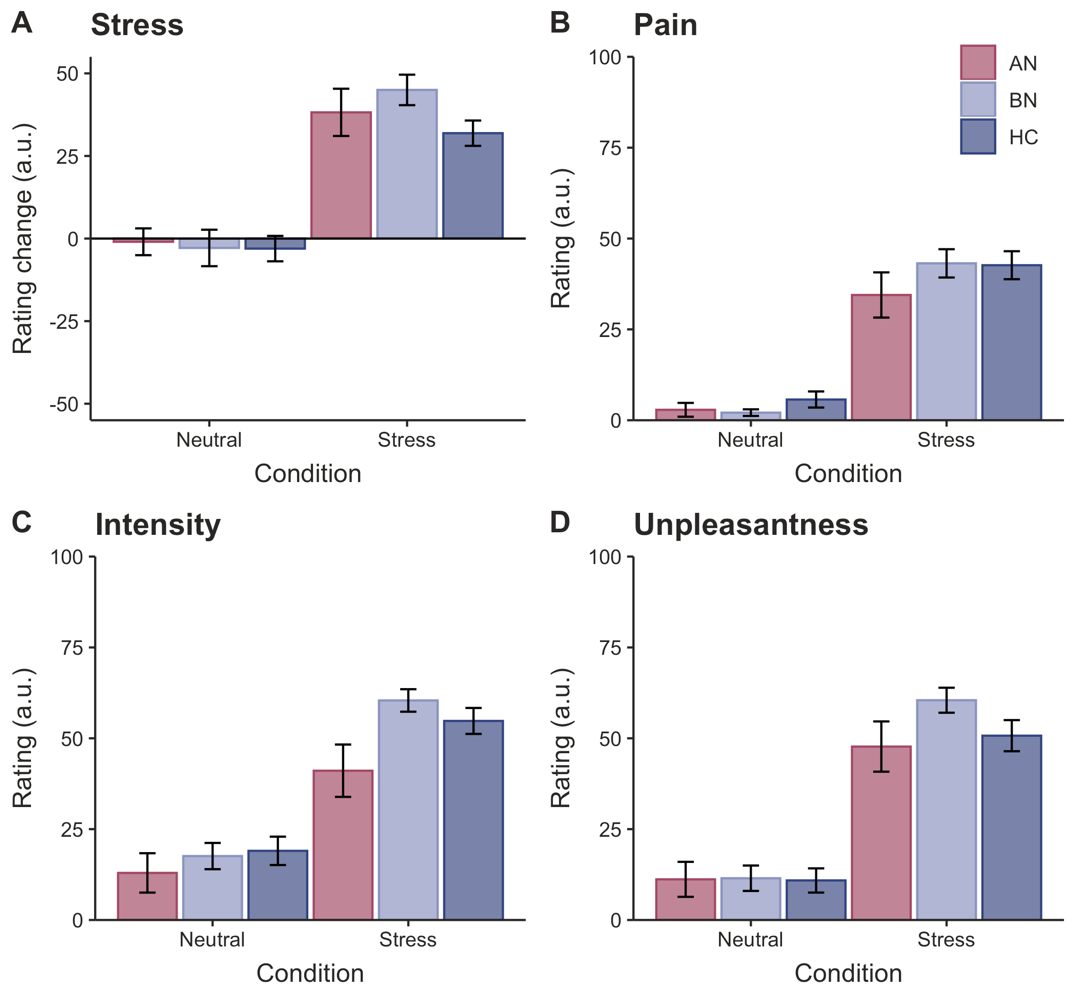

**Supplementary Figure 1.** *Participant ratings of induced stress and electrical stimulation.* **A)** The stress manipulation induced greater subjective stress compared to the neutral task. Participants rated the electrical stimulation as more **B)** painful, **C)** intense and **D)** unpleasant following stress as compared to the control task, where stimulation was intended to be detectable but not unpleasant. Ratings did not differ significantly by group (all p’s >.05). Error bars = SEM.

***1.3 Image acquisition***

MR scanning was completed at the Wolfson Brain Imaging Centre at Addenbrooke’s hospital on a 3T Siemens SkyraFit scanner (Erlangen, Germany). On each day, 1.0mm isotropic T1-weighted structural images were acquired (TE=2.95ms, TR=2300ms, flip angle=9°, acquisition matrix=256 X 256mm). Echo-planar images were acquired across 30 interleaved slices with the following parameters: TR=1600ms, TE=23ms, flip angle=78°, acquisition matrix=64x64, 3.0 mm isotropic voxels, 631 volumes.

***1.4 Functional MRI pre-processing:***

Anatomical scans were co-registered with a linear transformation (AFNI program *3dAllineate*) and averaged across days via *3dMean.* The averaged structural image was then processed with the standard FreeSurfer recon-all pipeline. The resulting white matter and ventricle segmentations were resampled to 3mm isotropic resolution and eroded by 1 voxel along each axis. Remaining pre-processing steps were completed with the afni_proc.py python script, in which functional images were slice-time corrected, re-aligned to the minimum outlier functional volume, co-registered to the subject’s skull-stripped averaged anatomical image, nonlinearly warped to the MNI152_T1_2009c template and smoothed using a 6mm full-width at half-maximum (FWHM) kernel. The first three principle components from the time series of lateral, third and fourth ventricle sources were estimated and regressed from functional volumes, along with six head motion parameters and their first-order derivatives. Local white matter was regressed from functional volumes using the fast *ANATICOR* pipeline (8). Functional volumes with a Euclidean norm motion derivative >0.5mm were censored, and participants with >10% of volumes censored were excluded from group-level analysis.

Functional MRI data from pre-stress, post-stress, pre-neutral and post-neutral sessions were available from n=84, n=79, n=80 and n=81 participants, respectively. One participant was excluded from analysis due to white matter abnormalities. In addition, 5 post-stress, 4 pre-neutral and 2 post-neutral runs were excluded because of excessive head motion. A technical error resulted in the exclusion of one additional post-neutral session. During a pre-neutral session, EPI acquisition had to be stopped due to a technical error; however, as ~70% of functional volumes had been acquired for this participant, their pre-neutral run was included in the group-level analysis.

***1.5 Regions of interest:***

A priori ROI selection was based on findings from previous functional imaging studies of the SSAT (9, 10), proactive and reactive inhibitory control networks (11), and NeuroSynth (<https://neurosynth.org>) clusters associated with “stop signal” and “response inhibition” terms. Anatomical ROIs were defined using the Brainnetome atlas (12), and average ROI beta estimates were extracted for each subject using *3dmaskave.*

***1.6 Exploratory analyses***

We used exploratory LMMs to examine whether brain regions demonstrating differing neural responses between groups (e.g., left premotor cortex, right IFG, left SFG) or in a group-by-condition-by-time interaction (right SFG, right vmPFC) explained variance in food intake (Z-scored). As described previously (1), one participant declined to initiate the ad libitum meal on Day 2, and another reported severe nausea prior to the meal, so we modelled observations from 83 participants for these exploratory analyses. For consistency, SSRT and neural responses were modelled from post-manipulation runs only.

Each model included fixed effects of group, condition and impulsivity measure, where random intercepts for within-subject variables were included within the subject’s random effect. Models of SSRT and brain responses also included a person-centered random slope for these variables. Group differences were assessed via nonorthogonal planned contrasts, comparing each patient group to controls.

2. **Supplementary Results**

***2.1 ROI analyses of reactive inhibition***

A significant main effect of time was related to right pre-supplementary motor cortex (β=-0.02, t(156)=-3.51, p<.001), anterior cingulate cortex (β=-0.01, t(156)=-2.79, p=.006) and bilateral superior parietal cortex activity (β=-0.02, t(156)=-4.46, p<.001) on Successful Stop versus Go trials, where activity declined post-manipulation. Moreover, the main effects of condition (β=0.01, t(82)=2.77, p=.007) and time (β=0.01, t(156)=3.14, p=.002) were associated with ACC activity during Successful vs Failed Stop trials, where *deactivation* was less negative on the stress day and post-runs. Interaction effects were nonsignificant across all ROIs for both contrasts. As a time-by-condition interaction term was not significantly related to ACC activity, the observed differences likely reflect BOLD variability across scan days that was not specific to the stress induction.

**Supplementary Table 1.** *Psychotropic medication by patient group*

| Medication | AN-BP | | BN | |
| --- | --- | --- | --- | --- |
|  | **# (%)** | **Dose (mg)** | **# (%)** | **Dose (mg)** |
| Amitriptyline | 1 (4.5) | 20 | - | - |
| Aripiprazole | 1 (4.5) | 5 | - | - |
| Bupropion | 1 (4.5) | 300 | - | - |
| Duloxetine | 1 (4.5) | 60 | - | - |
| Fluoxetine | 3 (13.6) | 40 – 60 | 4 (12.1) | 30 - 60 |
| Mirtazapine | - | - | 1 (3.0) | 15 |
| Olanzapine | 1 (4.5) | 5 | - | - |
| Sertraline | 2 (9.1) | 20 – 150 | 3 (9.1) | 100 |
| Venlafaxine | 1 (4.5) | 112.5 | 3 (9.1) | 150 – 300 |
| Zopiclone | - | - | 1 (3.0) | 3.75 |

**Note:** Several participants were prescribed more than one medication. Dose indicates mg/day.

**Supplementary Table 2.** SSAT performance metrics by group and condition

|  |  | **Neutral** | | **Stress** | |
| --- | --- | --- | --- | --- | --- |
|  |  | **Pre** | **Post** | **Pre** | **Post** |
| **Measure** | **Group** | **M ± 95% CI** | **M ± 95% CI** | **M ± 95% CI** | **M ± 95% CI** |
| *SSRT (ms)* | AN | 273 ± 4 | 269 ± 7 | 277 ± 5 | 268 ± 6 |
|  | BN | 270 ± 5 | 268 ± 5 | 269 ± 5 | 269 ± 5 |
|  | HC | 269 ± 6 | 268 ± 4 | 271 ± 5 | 267 ± 4 |
| *Go Trial 0%* | AN | 814.4 ± 1.1 | 808.7 ± 1.1 | 818.5 ± 1.2 | 813.2 ± 1.1 |
| (*ms)* | BN | 820.7 ± 1.0 | 815.0 ± 0.9 | 821.0 ± 1.0 | 816.7 ± 0.9 |
|  | HC | 823.6 ± 1.1 | 817.7 ± 1.0 | 818.9 ± 1.1 | 814.4 ± 1.0 |
|  |  | **M (SD)** | **M (SD)** | **M (SD)** | **M (SD)** |
| *Stop accuracy* | AN | 58.7 (4.1) | 57.7 (4.7) | 58.6 (4.2) | 58.8 (3.7) |
| *(%)* | BN | 60.2 (5.2) | 59.3 (6.0) | 60.1 (5.2) | 58.6 (4.2) |
|  | HC | 59.5 (4.7) | 59.1 (5.4) | 57.5 (5.1) | 57.7 (4.7) |
| *Accuracy* | AN | 98.3 (3.8) | 99.1 (0.9) | 98.0 (1.9) | 99.2 (0.9) |
| *(%)* | BN | 98.7 (1.3) | 98.9 (1.9) | 97.4 (5.7) | 99.4 (0.7) |
|  | HC | 98.0 (3.3) | 99.3 (0.9) | 98.4 (1.7) | 99.3 (0.9) |

Note: ‘Accuracy’ represents the percentage of go-signal trials on which participants made a response.

**Supplementary Table 3.** Linear mixed-effects analysis of reactive inhibition (Successful Stop > Go 0% contrast)

| Effect | Direction | Side | Region | Peak MNI Coordinates | | | Size (voxels) | F-statistic | Z-statistic |
| --- | --- | --- | --- | --- | --- | --- | --- | --- | --- |
|  |  |  |  | X | Y | Z |  |  |  |
| Time | Post < Pre | L | Superior medial gyrus, extending to SMA | 1 | 24 | 60 | 341 | 40.33 | -6.45 |
|  | Post < Pre | R | Middle frontal gyrus | 43 | 12 | 57 | 310 | 30.47 | -5.52 |
|  | Post < Pre | R | Angular gyrus | 52 | -57 | 39 | 166 | 30.97 | -5.56 |
|  | Post < Pre | R | Inferior frontal gyrus (pars opercularis) | 43 | 18 | -6 | 142 | 32.07 | -5.66 |
|  | Post > Pre | L | Primary sensory cortex | -41 | -42 | 66 | 120 | 21.55 | 4.64 |
|  | Post < Pre | L | Inferior frontal gyrus (pars opercularis) | -35 | 15 | -9 | 68 | 39.00 | -6.25 |
|  | Post < Pre | L | Cerebellum | -26 | -84 | -24 | 63 | 20.14 | -4.49 |
|  | Post < Pre | R | Superior frontal gyrus | 25 | 54 | 15 | 63 | 41.72 | -6.46 |
|  | Post < Pre | R | Middle temporal gyrus | 64 | -45 | 0 | 51 | 24.26 | -4.93 |
|  | Post > Pre | R | Superior parietal lobule | 16 | -57 | 75 | 30 | 17.98 | 4.24 |
|  | Post < Pre | L | Inferior temporal gyrus | -47 | -60 | -9 | 24 | 18.48 | -4.30 |
|  | Post < Pre | L | Precentral gyrus | -56 | 6 | 45 | 25 | 20.13 | -4.49 |
|  | Post < Pre | R | Caudate nucleus | 4 | -3 | 12 | 25 | 23.93 | -4.89 |
|  | Post < Pre | L | Premotor cortex | -44 | -3 | 45 | 25 | 17.48 | -4.18 |
|  | Post > Pre | R | Precentral gyrus | 37 | -24 | 72 | 21 | 17.28 | 4.16 |
|  | Post > Pre | L | Postcentral gyrus | -53 | -21 | 60 | 19 | 17.80 | 4.22 |

Note: Clusters were defined at a voxel-wise threshold of p < .001 and FWE-corrected at p < .05 (kE = 18.8 voxels). Cluster size was determined from the F-statistic map.

**Supplementary Table 4.** Whole-brain fMRI responses to stop-signal probability by group

| Group | Side | Region | Peak MNI Coordinates | | | Size (voxels) | Z-statistic |
| --- | --- | --- | --- | --- | --- | --- | --- |
|  |  |  | X | Y | Z |  |  |
| AN | R | Inferior occipital gyrus | 34 | -81 | -6 | 511 | 7.25 |
|  | L | Inferior occipital gyrus | -35 | -96 | -6 | 461 | 7.13 |
|  | R | Middle insula | 40 | 15 | 6 | 174 | 6.03 |
|  | L | Supramarginal gyrus | -62 | -30 | 45 | 109 | 5.03 |
|  | R | Superior occipital gyrus | 22 | -78 | 45 | 71 | 4.58 |
|  | R | Precentral gyrus | 52 | 9 | 33 | 69 | 4.53 |
|  | R | Supramarginal gyrus | 61 | -33 | 51 | 39 | 4.54 |
|  | L | Middle insula | -35 | 18 | 12 | 27 | 4.88 |
|  | R | Cerebellar vermis | 4 | -72 | -9 | 24 | 4.06 |
|  | L | Middle frontal gyrus | -26 | -3 | 60 | 21 | 4.39 |
| BN | L | Inferior occipital gyrus | -35 | -99 | -9 | 365 | 6.78 |
|  | R | Inferior occipital gyrus | 34 | -93 | -6 | 380 | 8.01 |
|  | L | Postcentral gyrus | -20 | -45 | 75 | 19 | -4.13 |
| HC | L | Fusiform gyrus | -35 | -78 | -9 | 639 | 7.77 |
|  | R | Inferior occipital gyrus | 31 | -90 | -6 | 441 | 9.28 |
|  | R | Intraparietal sulcus | 31 | -78 | 45 | 94 | 4.61 |
|  | R | Precentral gyrus | 46 | 9 | 33 | 60 | 5.08 |
|  | L | Rostral middle frontal gyrus | -20 | 57 | 36 | 52 | -4.35 |
|  | L | Inferior frontal gyrus (pars orbitalis) | -44 | 27 | -9 | 51 | -5.32 |
|  | L | Middle frontal gyrus | -44 | 24 | 51 | 45 | -4.58 |
|  | R | Postcentral gyrus | 61 | -24 | 39 | 31 | 5.17 |
|  | L | Supramarginal gyrus | -62 | -30 | 39 | 28 | 4.35 |
|  | R | SMA | 4 | 6 | 60 | 24 | 4.96 |
|  | L | Superior frontal gyrus | -5 | 42 | 57 | 23 | -4.17 |

Note: Clusters were defined at a voxel-wise threshold of p < .001 and FWE-corrected at p < .05 (kE = 18.8 voxels).

**Supplementary Table 5.** Whole-brain responses to reaction time amplitude modulator by group

| Group | Side | Region | Peak MNI Coordinates | | | Size (voxels) | Z-statistic |
| --- | --- | --- | --- | --- | --- | --- | --- |
|  |  |  | X | Y | Z |  |  |
| AN | L | Primary motor cortex | -41 | -27 | 60 | 1846 | 7.80 |
|  | R | Inferior occipital gyrus | 34 | -93 | 3 | 561 | 6.97 |
|  | R | Superior parietal lobule | 16 | -60 | 72 | 352 | 6.26 |
|  | R | SMA | 1 | -9 | 57 | 336 | 7.84 |
|  | R | Cerebellar vermis | 4 | -72 | -30 | 300 | 6.20 |
|  | R | Superior frontal gyrus | 25 | -12 | 69 | 170 | 7.11 |
|  | L | Anterior cingulate cortex | -14 | 42 | 12 | 157 | -4.98 |
|  | L | Calcarine sulcus | -14 | -81 | 15 | 142 | 5.34 |
|  | R | Supramarginal gyrus | 67 | -33 | 33 | 108 | 4.87 |
|  | L | Middle cingulate cortex | -2 | -39 | 45 | 108 | -4.86 |
|  | L | Angular gyrus | -44 | -72 | 57 | 50 | -4.39 |
|  | L | Supramarginal gyrus | -50 | -30 | 21 | 41 | 5.00 |
|  | L | Thalamus (prefrontal) | -14 | -24 | 12 | 30 | 4.54 |
|  | L | Middle frontal gyrus | -38 | 6 | 63 | 29 | -4.28 |
|  | R | Frontal opercular | 49 | 0 | 12 | 24 | 4.57 |
|  | R | Cerebellar cortex | 16 | -96 | -30 | 19 | -4.98 |
|  | R | Caudate nucleus | 7 | 6 | 0 | 19 | -3.90 |
| BN | L | Precuneus | -11 | -66 | 69 | 1688 | 8.65 |
|  | R | Precuneus | 10 | -60 | 72 | 818 | 8.74 |
|  | L | SMA | -2 | -6 | 60 | 382 | 7.50 |
|  | R | Medial frontal gyrus | 7 | 30 | -9 | 224 | -5.37 |
|  | R | Cerebellar cortex | 19 | -63 | -18 | 222 | 4.97 |
|  | R | Middle temporal gyrus | 52 | -66 | 25 | 199 | 4.93 |
|  | R | Superior frontal gyrus | 28 | -9 | 72 | 184 | 5.86 |
|  | L | Middle occipital gyrus | -41 | -81 | 45 | 172 | -5.14 |
|  | R | Supramarginal gyrus | 61 | -36 | 45 | 147 | 4.70 |
|  | L | Caudate nucleus | -5 | 6 | 3 | 126 | -5.54 |
|  | L | Inferior frontal gyrus (pars orbitalis) | -53 | 24 | 0 | 111 | -4.82 |
|  | L | Superior frontal gyrus | -11 | 57 | 42 | 85 | -4.51 |
|  | L | Middle occipital gyrus | -50 | -87 | 6 | 75 | 5.00 |
|  | L | Middle cingulate cortex | -5 | -42 | 45 | 73 | -5.04 |
|  | L | Middle frontal gyrus | -23 | 18 | 57 | 71 | -4.86 |
|  | R | Cerebellar cortex | 46 | -75 | -30 | 67 | -6.59 |
|  | L | Lingual gyrus | -23 | -96 | -9 | 64 | 4.80 |
|  | L | Cerebellar cortex | -35 | -48 | -27 | 62 | 5.09 |
|  | R | Cerebellar cortex | 10 | -87 | -21 | 54 | -5.03 |
|  | L | Cuneus | -20 | -78 | 9 | 34 | 4.88 |
|  | L | Parieto-occipital sulcus | -11 | -60 | 21 | 30 | -5.04 |
|  | L | Superior frontal gyrus | -23 | 57 | 9 | 29 | -4.56 |
|  | L | Orbitofrontal cortex | -26 | 21 | -15 | 27 | -4.74 |
|  | L | Rostral middle frontal | -56 | 18 | 36 | 24 | -4.66 |
|  | L | Caudal middle frontal | -41 | 12 | 39 | 21 | -4.68 |
|  | L | Inferior temporal gyrus | -56 | -51 | -9 | 20 | -5.34 |
| HC | L | Precentral gyrus | -30 | -7 | 66 | 2854 | 9.21 |
|  | R | Precuneus | 7 | -63 | 72 | 1601 | 8.43 |
|  | R | Cerebellar vermis (VII) | 4 | -78 | -27 | 471 | 6.75 |
|  | R | Anterior cingulate cortex | 4 | 36 | 6 | 191 | -5.56 |
|  | L | Middle occipital gyrus | -38 | -93 | 0 | 130 | 4.66 |
|  | L | Cerebellar vermis (VI) | -35 | -48 | -27 | 111 | 5.66 |
|  | L | Putamen | -14 | -3 | -6 | 90 | -4.69 |
|  | L | Middle orbital gyrus | -5 | 54 | 3 | 75 | -5.38 |
|  | L | Thalamus (prefrontal) | -14 | -24 | 12 | 64 | 7.23 |
|  | R | Thalamus (prefrontal) | 10 | -21 | 12 | 63 | 6.09 |
|  | R | Inferior occipital gyrus | 22 | -93 | -3 | 55 | 5.56 |
|  | L | Calcarine sulcus | -11 | -60 | 12 | 33 | -4.74 |
|  | R | Thalamus/red nucleus | 7 | -24 | -6 | 22 | 4.83 |
|  | R | Fusiform gyrus | 25 | -87 | -12 | 21 | 4.75 |
|  | L | Medial frontal gyrus | -2 | 57 | 21 | 20 | -4.16 |
|  | L | Angular gyrus | -38 | -78 | 48 | 20 | -3.90 |
|  | R | Prostriate area | 28 | -54 | 18 | 19 | -4.14 |

Note: Clusters were defined at a voxel-wise threshold of p < .001 and FWE-corrected at p < .05 (kE = 18.8 voxels).

**Supplementary Table 6.** Whole-brain responses during reactive inhibition (Successful Stop > Go-signal 0%) by group

| Group | Side | Region | Peak MNI Coordinates | | | Size (voxels) | Z-statistic |
| --- | --- | --- | --- | --- | --- | --- | --- |
|  |  |  | X | Y | Z |  |  |
| AN | L | Postcentral gyrus | -32 | -36 | 69 | 6396 | -10.77 |
|  | R | Inferior frontal gyrus (pars opercularis) | 40 | 12 | 9 | 3914 | 10.98 |
|  | R | Medial orbitofrontal | 7 | 33 | -9 | 2828 | -9.11 |
|  | R | Middle occipital gyrus | 31 | -72 | 39 | 1951 | 8.03 |
|  | L | Middle occipital gyrus | -44 | -81 | 42 | 335 | -6.82 |
|  | L | Inferior parietal lobule | -29 | -63 | 54 | 250 | 7.28 |
|  | L | Fusiform gyrus | -35 | -63 | -9 | 217 | 6.15 |
|  | L | Cerebellar cortex (VI) | -35 | -57 | -30 | 200 | 7.30 |
|  | R | Cerebellar cortex (VIIa) | 34 | -84 | -30 | 190 | -6.91 |
|  | R | Superior frontal gyrus | 22 | 30 | 54 | 172 | -7.69 |
|  | L | Rostral middle frontal gyrus | -38 | 42 | 36 | 116 | 5.69 |
|  | R | Angular gyrus | 49 | -72 | 39 | 114 | -6.12 |
|  | R | Middle cingulate cortex | 7 | -24 | 33 | 81 | 5.89 |
|  | L | Supramarginal gyrus | -65 | -33 | 30 | 74 | 6.21 |
|  | R | Inferior occipital gyrus | 25 | -105 | 3 | 70 | -6.67 |
|  | R | Superior temporal gyrus | 49 | -30 | 3 | 65 | 5.95 |
|  | L | Middle occipital gyrus | -29 | -102 | 0 | 56 | -5.74 |
|  | L | Cerebellar cortex (VIIa) | -17 | -93 | -33 | 45 | -5.29 |
|  | R | Lateral orbitofrontal | 37 | 30 | -12 | 40 | -6.98 |
|  | R | Cerebellar cortex (VI) | 34 | -57 | -27 | 36 | 6.02 |
|  | R | Caudate nucleus | 16 | 15 | 21 | 27 | -5.32 |
| BN | L | Calcarine sulcus | -8 | -69 | -21 | 6350 | -10.26 |
|  | R | Inferior occipital gyrus | 4 | 9 | 54 | 5381 | 12.34 |
|  | R | Intraparietal sulcus | 31 | -75 | 36 | 3430 | 10.52 |
|  | L | Medial orbitofrontal | -8 | 39 | -9 | 2572 | -10.26 |
|  | L | Cerebellar cortex (VI) | -32 | -60 | -27 | 1887 | 9.26 |
|  | L | Angular gyrus | -53 | -72 | 33 | 550 | -8.69 |
|  | R | Cerebellum | 16 | -93 | -33 | 227 | -8.85 |
|  | R | Superior frontal gyrus | 19 | 33 | 60 | 181 | -6.82 |
|  | L | Rostral middle frontal gyrus | -35 | 42 | 30 | 172 | 6.27 |
|  | R | Angular gyrus | 52 | -72 | 36 | 138 | -6.89 |
|  | R | Superior temporal gyrus | 49 | -27 | 0 | 92 | 6.44 |
|  | L | Cerebellar cortex (VIIa) | -17 | -93 | -33 | 85 | -6.89 |
|  | L | Middle occipital gyrus | -23 | -108 | 3 | 67 | -6.31 |
|  | R | Inferior occipital gyurs | 28 | -105 | 0 | 64 | -8.25 |
|  | R | Caudate | 19 | -18 | 36 | 57 | -6.21 |
|  | R | Prostriate area | 31 | -54 | 6 | 48 | -6.83 |
|  | R | Inferior frontal gyrus (pars orbitalis) | 37 | 30 | -12 | 37 | -6.12 |
|  | R | Dorsolateral putamen | 28 | -24 | 9 | 28 | 5.40 |
| HC | L | Calcarine sulcus | -8 | -69 | 21 | 11370 | -10.77 |
|  | R | Inferior frontal gyrus (pars opercularis) | 40 | 15 | 9 | 4124 | 12.03 |
|  | R | Middle occipital sulcus | 34 | -75 | 36 | 2264 | 9.69 |
|  | L | Insula | -32 | 15 | 9 | 714 | 12.14 |
|  | L | Superior parietal lobule | -26 | -69 | 57 | 563 | 7.13 |
|  | L | Fusiform gyrus | -35 | -63 | -6 | 508 | 8.53 |
|  | L | Middle cingulate cortex | -8 | -30 | 33 | 237 | 7.92 |
|  | L | Cerebellar cortex (VIIa) | -17 | -93 | -33 | 194 | -8.09 |
|  | R | Superior frontal gyrus | 22 | 30 | 54 | 146 | -6.80 |
|  | L | Cerebellar cortex (VI) | -32 | -57 | -30 | 123 | 8.20 |
|  | R | Angular gyrus | 49 | -72 | 39 | 101 | -6.57 |
|  | L | Middle frontal gyrus | -35 | 42 | 30 | 100 | 6.38 |
|  | L | Cerebellar cortex | -8 | -78 | -18 | 75 | 6.28 |
|  | R | Superior temporal gyrus | 49 | -30 | 0 | 59 | 5.24 |
|  | R | Dorsal caudate | 16 | 0 | 33 | 42 | -6.21 |
|  | R | Cerebellar cortex (VI) | 34 | -57 | -27 | 28 | 5.47 |
|  | R | Inferior frontal gyrus (pars orbitalis) | 37 | 30 | -12 | 28 | -4.94 |
|  | R | Caudate | 19 | -18 | 36 | 28 | -6.11 |

Note: Clusters were defined at a voxel-wise threshold of p < .001 and FWE-corrected at p < .05 (kE = 18.8 voxels).

**Supplementary Table 7.** Whole-brain responses during reactive inhibition (Successful Stop > Failed Stop) by group

| Group | Side | Region | Peak MNI Coordinates | | | Size (voxels) | Z-statistic |
| --- | --- | --- | --- | --- | --- | --- | --- |
|  |  |  | X | Y | Z |  |  |
| AN | R | Cerebellum (IV-V) | 10 | -60 | -18 | 3659 | -9.24 |
|  | L | Postcentral gyrus | -44 | -24 | 57 | 1154 | -9.29 |
|  | R | Middle cingulate cortex | 1 | 18 | 39 | 1104 | -7.96 |
|  | L | Inferior frontal gyrus (pars opercularis) | -53 | 9 | 3 | 434 | -8.43 |
|  | R | Inferior frontal gyrus (pars opercularis) | 52 | 9 | 0 | 393 | -8.20 |
|  | R | SMA | 10 | 6 | 72 | 273 | -6.12 |
|  | R | Middle temporal gyrus | 58 | -48 | 15 | 141 | -4.62 |
|  | R | Rostral middle frontal | 28 | 54 | 30 | 124 | -5.08 |
|  | R | Middle temporal gyrus | 46 | -75 | 18 | 115 | -6.03 |
|  | L | Middle frontal gyrus | -32 | 51 | 27 | 109 | -4.87 |
|  | L | Superior parietal lobule | -32 | -69 | 63 | 70 | -4.85 |
|  | R | Cerebellar vermis | 4 | -75 | -30 | 45 | -6.85 |
|  | R | Primary sensory cortex | 19 | -33 | 69 | 43 | -4.61 |
|  | R | Precentral gyrus | 49 | 3 | 54 | 42 | -4.60 |
|  | R | Inferior parietal lobule | 49 | -51 | 60 | 29 | -4.91 |
|  | R | Superior parietal lobule | 22 | -78 | 57 | 26 | -4.06 |
|  | L | Posterior insula | -38 | -15 | -3 | 23 | -4.61 |
|  | R | Putamen | 28 | -6 | 6 | 22 | 4.24 |
|  | L | Middle temporal gyrus | -59 | -60 | 18 | 21 | -4.10 |
|  | L | Caudate nucleus | -8 | 18 | 6 | 20 | -4.33 |
|  | R | Postcentral gyrus | 64 | -18 | 33 | 20 | -4.14 |
|  | R | Precuneus | 10 | -75 | 66 | 20 | -4.87 |
| BN | L | Middle occipital gyrus | -32 | -87 | 24 | 12819 | -11.61 |
|  | R | Inferior occipital gyrus | 34 | -93 | -6 | 725 | -7.60 |
|  | L | Insula | -32 | 15 | -6 | 608 | -7.97 |
|  | R | Superior frontal gyrus | 22 | 51 | 30 | 164 | -6.54 |
|  | L | Middle frontal gyrus | -32 | 45 | 24 | 97 | -5.18 |
|  | R | Precentral gyrus | 25 | -33 | 78 | 83 | -5.07 |
|  | L | Putamen | -29 | -3 | 6 | 71 | 5.84 |
|  | R | Putamen | 25 | 3 | 0 | 48 | 5.27 |
|  | L | Angular gyrus | -50 | -72 | 54 | 43 | 5.73 |
|  | L | Middle frontal gyrus | -26 | 24 | 63 | 34 | 4.31 |
|  | R | Posterior insula | 40 | -15 | 3 | 31 | -4.67 |
|  | R | Inferior temporal gyrus | 46 | -6 | -27 | 30 | -4.71 |
|  | L | Middle temporal gyrus | -62 | -21 | 6 | 28 | 4.39 |
|  | L | Inferior frontal gyrus (pars orbitalis) | -41 | 42 | -6 | 24 | 4.75 |
| HC | L | Anterior cingulate cortex | -5 | 18 | 36 | 11517 | -10.05 |
|  | R | Inferior frontal gyrus (pars opercularis) | 49 | 9 | 6 | 402 | -8.04 |
|  | R | Precentral gyrus | 49 | 3 | 54 | 273 | -6.56 |
|  | L | Inferior parietal lobule | -50 | -66 | 54 | 50 | 5.12 |
|  | L | Middle temporal gyrus | 49 | -24 | -6 | 39 | -4.50 |
|  | L | Para-insular area | -41 | -18 | -3 | 29 | -5.85 |
|  | R | Middle frontal gyrus | 31 | 36 | 30 | 26 | -4.08 |
|  | L | Medial frontal gyrus | -8 | 33 | -9 | 23 | 5.39 |
|  | L | Middle frontal gyrus | -29 | 42 | 33 | 23 | -5.03 |
|  | L | Middle frontal gyrus | -32 | 21 | 57 | 22 | 4.11 |
|  | R | Superior frontal gyrus | 34 | 54 | 6 | 21 | 3.90 |

Note: Clusters were defined at a voxel-wise threshold of p < .001 and FWE-corrected at p < .05 (kE = 18.8 voxels).

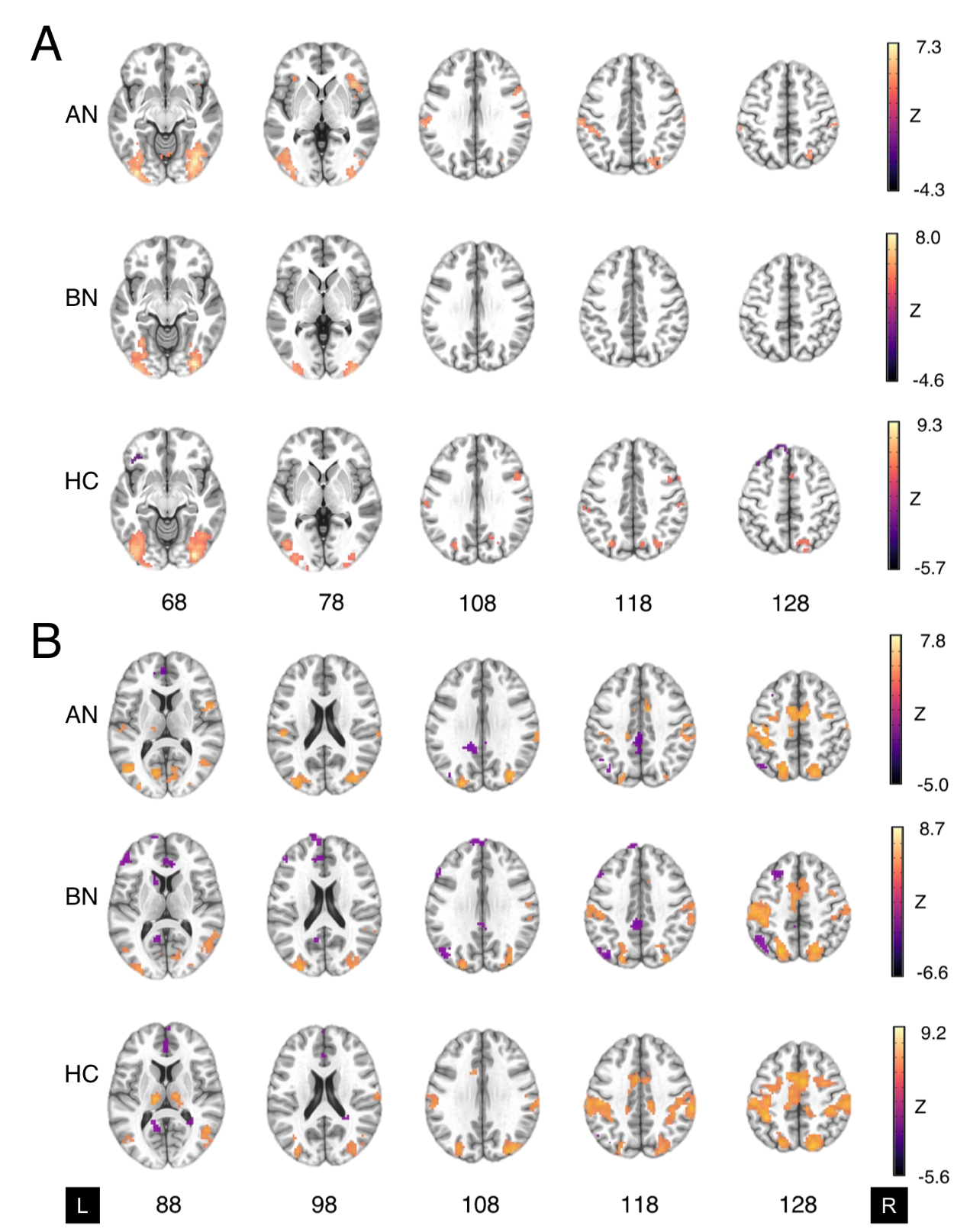

**Supplementary Figure 2.** *Whole-brain activation in anorexia nervosa, bulimia nervosa and controls during proactive inhibition.* Two-sample t-tests of **A)** the parametric effect of stop-signal probability versus the implicit baseline (i.e., Go_0%_ trials) and **B)** the parametric effect of reaction time versus the implicit baseline activation for AN-BP, BN and control groups. Maps represent significant clusters (voxel-wise p-value < .001, FWE cluster probability p-value < .05). and are presented in neurological orientation (L=left).

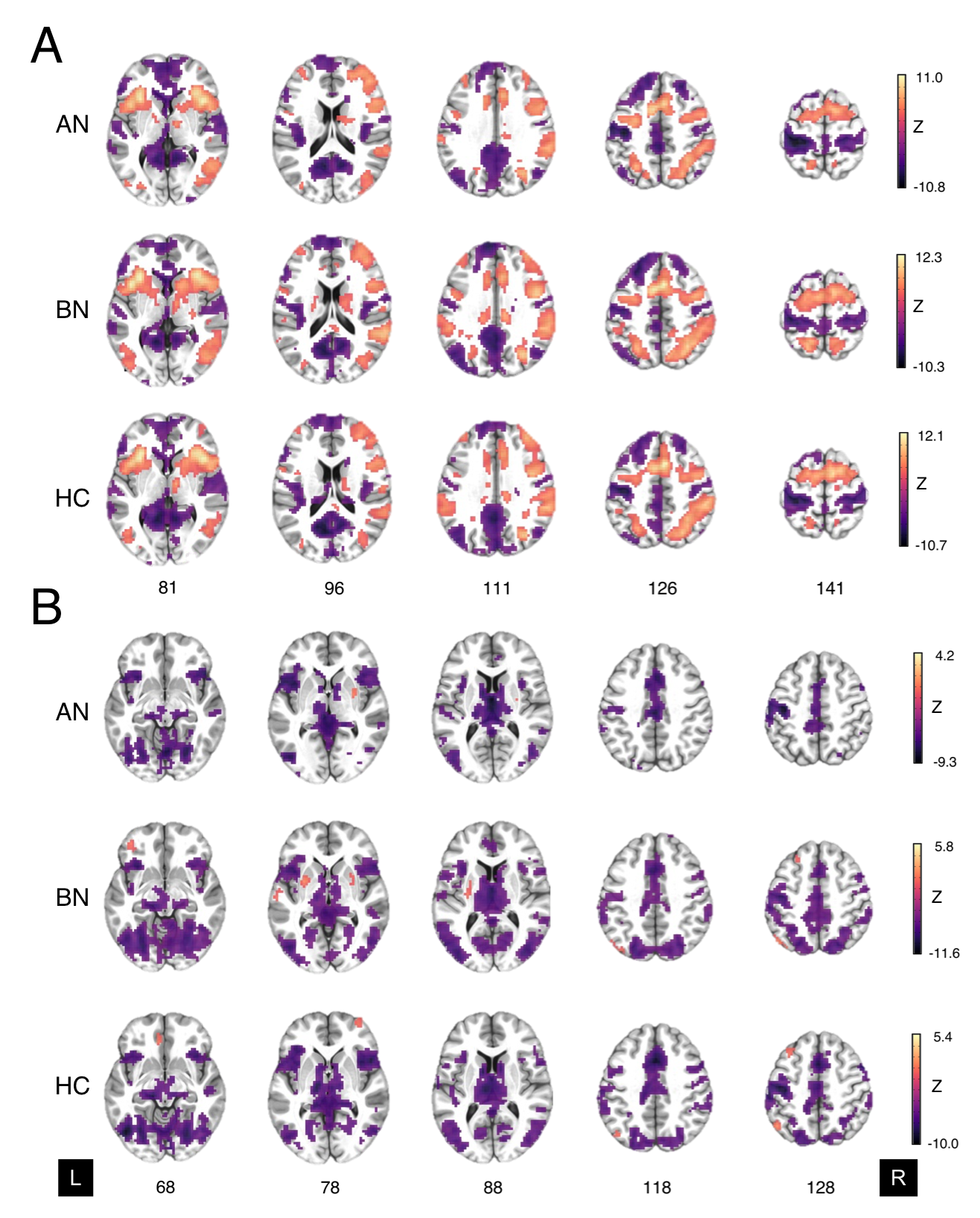

**Supplementary Figure 3.** *Whole-brain activation in anorexia nervosa, bulimia nervosa and controls during reactive inhibition.* Two-sample t-tests of **A)** successful stop-signal versus baseline Go trials and **B)** successful stop-signal versus failed stop-signal activation for AN-BP, BN and control groups. As contrasts were computed in the *3dLME* AFNI package, statistical maps represent Z-scores, as denoted by the color bar. Maps represent significant clusters (voxel-wise p-value < .001, FWE cluster-probability p-value < .05) and are presented in neurological orientation (L=left).

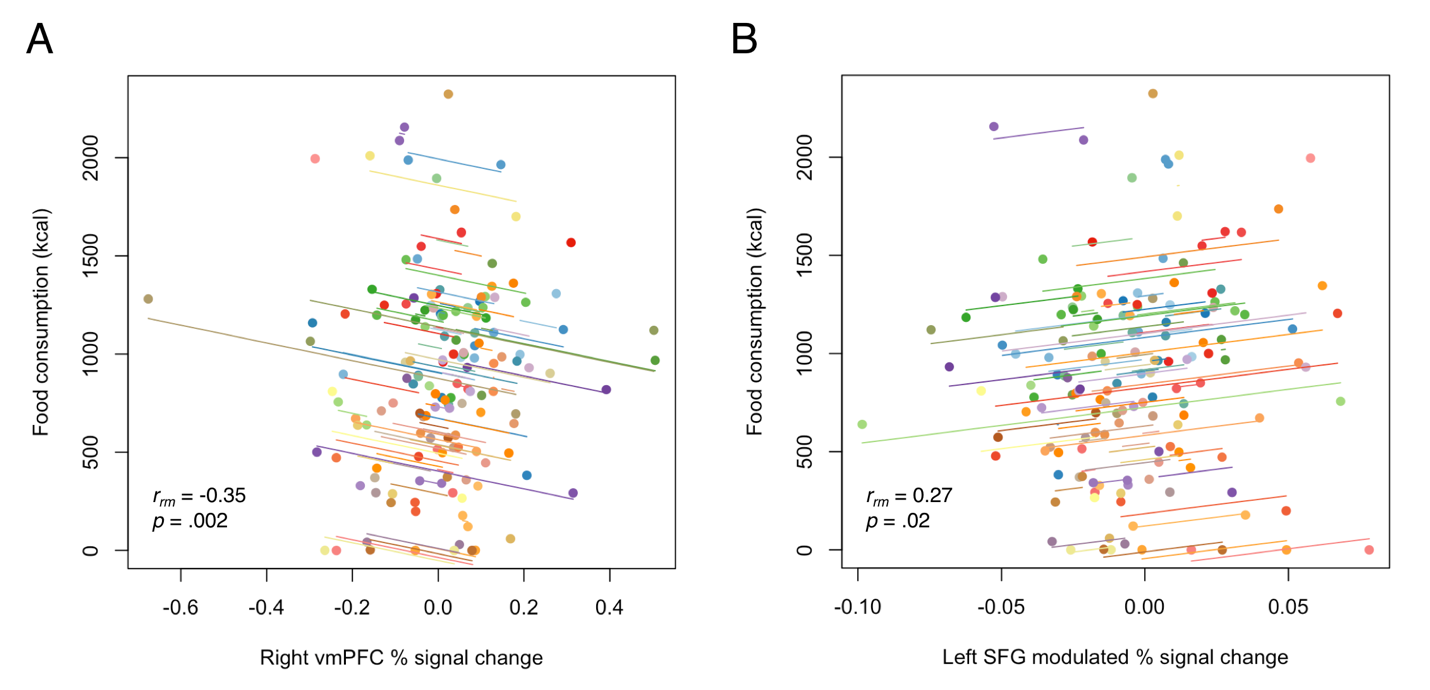

**Supplementary Figure 4.** *Associations between prefrontal responses during inhibition and ad libitum consumption.* **A)** Greater vmPFC responses during reactive inhibition (Successful Stop vs. Failed Stop) were negatively related to food consumption during the free choice meal. **B)** Increased left superior frontal gyrus responses to greater stop-signal probability were positively associated with food intake. Observations within the same subject are given the same colour with a corresponding line, representing the overall line of best fit. While effects were derived from linear mixed-effects models, repeated measures correlations were computed for visualisation, using the *rmcorr* R package (13) .
